# Shared genetic underpinnings between genetic generalized epilepsy and background EEG oscillations

**DOI:** 10.1101/711200

**Authors:** Remi Stevelink, Jurjen J. Luykx, Bochao D. Lin, Costin Leu, Dennis Lal, Alexander Smith, Dick Schijven, Johannes A. Carpay, Koen Rademaker, Roiza A. Rodrigues Baldez, Orrin Devinsky, Kees P.J. Braun, Floor E. Jansen, ILAE Consortium on Complex Epilepsies, Epi25 Collaborative, Dirk Smit, Bobby P.C. Koeleman

## Abstract

**Objective:** Paroxysmal epileptiform abnormalities on electroencephalography (EEG) are the hallmark of epilepsies, but it is uncertain to what extent epilepsy and background EEG oscillations share neurobiological underpinnings. Here, we aimed to assess the genetic correlation between epilepsy and background EEG oscillations.

**Methods:** Confounding factors, including the heterogeneous etiology of epilepsies and medication effects hamper studies on background brain activity in people with epilepsy. To overcome this limitation, we compared genetic data from a GWAS on epilepsy (n=12,803 people with epilepsy and 24,218 controls) with that from a GWAS on background EEG (n=8,425 subjects without epilepsy), in which background EEG oscillation power was quantified in four different frequency bands: alpha, beta, delta and theta. We replicated our findings in an independent epilepsy replication dataset (n=4851 people with epilepsy and 20,428 controls). To assess the genetic overlap between these phenotypes, we performed genetic correlation analyses using linkage disequilibrium score regression, polygenic risk scores and Mendelian randomization analyses.

**Results:** Our analyses show strong genetic correlations between genetic generalized epilepsy (GGE) with background EEG oscillations, primarily with the beta frequency band. Furthermore, we show that subjects with higher beta and theta polygenic risk scores have a significantly higher risk of having generalized epilepsy. Mendelian randomization analyses suggest a causal effect of GGE genetic liability on beta oscillations.

**Significance:** Our results point to shared biological mechanisms underlying background EEG oscillations and the susceptibility for GGE, opening avenues to investigate the clinical utility of background EEG oscillations in the diagnostic work-up of epilepsy.

## Introduction

The power of oscillations in background EEG is a highly stable and heritable human trait ^1^. It is easily acquired and can be automatically analyzed by software, rather than subjective interpretation. Epilepsy is highly heritable and is characterized by altered brain excitability ^2, 3^. Oscillatory activity is believed to serve an essential role in cortico-thalamic functioning, and can be measured as power of oscillations in background EEG at different broad-band frequencies ^4^. Neurophysiological relationships between background EEG and generalized epileptiform discharges have been well described ^5–8^. However, it is currently unknown whether background oscillatory activity is itself associated with epilepsy, and whether background EEG and epilepsy have a shared neurobiological and genetic basis.

There have been some studies where background EEG oscillations measurements have been directly compared between people with epilepsy and healthy controls. However, such studies have yielded conflicting results, most likely because sample sizes were small and antiseizure drugs can strongly affect EEG measurements ^11–16^. These limitations and bias can be overcome by large-scale genetic studies, in which genetic determinants of background EEG measurements are assessed independently in healthy controls (presumably not taking antiseizure drugs). These genetic determinants can than be compared to genetic determinants of different epilepsy phenotypes, as assessed in a different study. Comparing these independent studies allows for a well-powered and unbiased assessment of shared genetic determinants of epilepsy and EEG oscillations.

Here, we therefore assessed whether oscillatory background EEG is genetically correlated with focal and generalized epilepsy. The association between genetic variants and background brain activity was previously investigated in a genome-wide association study (GWAS) on 8,425 subjects without epilepsy ^9^ We combined this data with our recently published large GWAS of epilepsy ^10^, to examine genetic correlations between several types of epilepsy and oscillatory brain activity across frequency bands (delta: 1-3.75Hz, theta: 47–75 Hz, alpha: 12.75 Hz and beta: 13-30Hz). Next, we utilized polygenic risk scoring (PRS) to assess whether patients with GGE have a genetic predisposition towards altered background brain activity. We then replicated genetic correlation and polygenic analyses using an independent cohort from the Epi25 Collaborative (n=4,851 people with epilepsy and 20,428 controls). Finally, we performed Mendelian Randomization to gain insight into possible causal relationships between genetic variants associated with epilepsy and those associated with background EEG. We thus provide converging evidence for consistent cross-trait genetic overlap between epilepsy and background EEG.

## Methods

### Study population: discovery dataset

The participants derived from the epilepsy GWASs ^10^ for the current analyses were Caucasian subjects. The epilepsy GWAS included 13 control cohorts ^10^. Case/control ascertainment and diagnostic criteria were previously reported ^10^. As described previously ^10^, epilepsy specialists diagnosed patients and ascertained phenotypic subtypes. Population-based datasets, some of which had been screened to exclude neurological disorders, were used as controls. However, due to the relatively low prevalence of epilepsy in the general population (~0.5-1%), screening to exclude epilepsy in control cohorts it will only have a minor effect on statistical power. Summary statistics from the recent epilepsy GWAS conducted by the ILAE Consortium on Complex Epilepsies GWAS were available for n=12,803 cases (suffering from either focal or generalized epilepsy) and 24,218 controls ^10^. From those participants the following subjects were excluded for those analyses requiring individual-level genotype data: Finnish ancestry (none had GGE) and the subset of the EPICURE-SP1 cohort that lacked informed consent for the current analyses, resulting in subject-level genotype data being available for 11,446 patients with epilepsy and 22,078 controls. Subjects with epilepsy were stratified into GGE (n=3,122) and focal epilepsy (n=8,324); GGE was further subdivided into childhood absence epilepsy (CAE; n=561), juvenile absence epilepsy (JAE; n=311), juvenile myoclonic epilepsy (JME; n=1,000) and generalized tonic-clonic seizures only (GTCS only; n=195). GGE subtype information was not available for 1055 patients.

We downloaded summary statistics of the ENIGMA-EEG GWAS of resting state oscillation power in the delta (1-3 Hz), theta (4-8 Hz), alpha (8-12 Hz), and beta (13-30 Hz) bands at the vertex (Cz) electrode (n=8,425 participants) ^9^ This EEG GWAS study was based on 5 cohorts from 4 cooperating centers. Although the selection criteria varied across cohorts, all adult cohorts included epilepsy and prolonged unconsciousness after head trauma as exclusion criteria which were communicated at the time of recruitment or at the first lab visit; since neurological disorders were an exclusion criterium, we do not expect subjects to be taking anti-seizure drugs (although this was no explicit exclusion criterium). All these were self or parent-reported retrospective questions. A fuller sample description is presented in the original study ^9^.

Approval for the source studies was obtained by all relevant institutional review boards and all study participants provided written informed consent according to the Declaration of Helsinki.

### Replication dataset

To replicate our findings, we used data from the Epi25 collaborative (http://epi-25.org/). This cohort currently constitutes 4,851 people with epilepsy, of whom 2,612 have focal epilepsy and 2,239 have GGE (no data on GGE subtypes were available). The cases were matched to a total of 20,428 controls from the Partners Healthcare Biobank (n=14,857), the Epi25 collaborative (n=210), the Genetics and Personality consortium (n=456), and an in-house project on inflammatory bowel disease (n=4,905). The cohorts were genotyped on the Illumina Global Screening Array, with the exception of the Partners Healthcare Biobank participants, who were genotyped on the Illumina Multi-Ethnic Screening Array. Approval was obtained by all relevant institutional review boards and all study participants provided written informed consent according to the Declaration of Helsinki.

### Genetic correlation analyses

Genetic correlations between epilepsy subtypes and oscillatory brain activity were computed using bivariate linkage-disequilibrium score regression (LDSC) ^22^ For these analyses, as no individual-level genotype data were available from the EEG dataset, we used published summary statistics of the EEG frequency bands (alpha, beta, delta and theta; n=8,425 participants) and the epilepsy subtypes (focal, GGE, CAE, JAE, JME and GTCS only; n=12,803 cases suffering from either focal or generalized epilepsy and 24,218 controls) from the ILAE consortium as a discovery dataset ^10^. For LDSC replication analyses, we used unpublished data from the Epi25 collaborative (http://epi-25.org/; n=4,851 people with epilepsy and 20,428 controls). For discovery and replication LDSC analyses, default settings of LDSC were used, with pre-computed LD score weights derived from the European subset of the 1000 Genomes project ^23^. See Supplemental Table 1 for the numbers of SNPs per LDSC analysis. The significance threshold was Bonferroni-corrected for the two main epilepsy subtypes studied (GGE and focal) but not for the EEG power spectra, since these were all highly correlated at p<10^-17^ (Supplemental Table 2), resulting in a significance threshold of p=0.05/2=0.025. Similarly, we did not correct for the individual GGE subtypes, which are phenotypically similar and genetically highly correlated ^10^.

### Polygenic risk score analyses

For PRS analyses we used individual-level genotype data derived from the epilepsy GWAS ^10^ and summary statistics from the EEG GWAS ^9^ Quality control was performed as reported in the published epilepsy GWAS ^10^. We then added a genotype filter for call rate >0.99 and the exclusion of genetically related subjects to allow for highly conservative PRS estimates. Genetic inter-relatedness was calculated with KING ^24^ and one subject from each pair with third-degree or higher relatedness (kinship coefficient >0.0442) was excluded. PRSice ^25^ was used with default settings to assess whether subjects with epilepsy have different EEG frequency power PRSs compared to controls. In brief, to each SNP we assigned a weight proportional to its association in the four EEG GWASs (alpha, beta, delta and theta). Next, individual PRSs were calculated as the sum of weighted effect alleles for every subject from the epilepsy cohort. These PRSs were standardized with a Z-score transformation 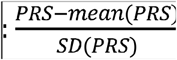. SNPs were pruned to a subset of genetically uncorrelated SNPs (linkage disequilibrium R^2^<0.1) and PRS values were calculated using a number of different P-value thresholds (P_t_s) from 0.0001 to 0.5. Next, logistic regression analyses, corrected for sex and 10 genetic ancestry principal components, were performed to assess the association of these PRS scores with GGE. The PRSs with the highest association with GGE was chosen as the ‘best-fit’, after which logistic regression analyses were repeated to assess the association of this PRS with the other epilepsy subtypes. We used a conservative p<0.001 significance threshold to correct for multiple comparisons, as recommended for PRSice ^25^. Δ Explained variance represented by the Nagelkerke R^2^ was computed using a logistic regression of the PRS, subtracted by the baseline model (covariates only: sex and 4 PCs). To quantify the association of beta-power PRS with GGE, we used PRSice standard settings to divide subjects into 10 deciles based on their beta-power PRS scores. We then performed logistic regression to compare the risk of having GGE between every decile with the lowest (0-10%) as a reference (corrected for sec and 4 PCs). We then repeated the analyses in the independent Epi25 cohort. This dataset contained approximately one-third fewer GGE cases than the discovery cohort, providing insufficient power to exactly replicate our discovery PRS findings. We therefore performed quasi-replication using a one-sample test of the proportion to assess concordance effect directions between discovery and replication PRS analyses, computing z-scores that were converted into p-values:

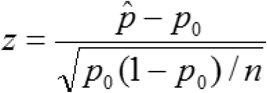

where the p = the sample proportion; H_0_ represents the null hypothesis: p = p_0_; and the alternative hypothesis H_1_: p ≠ p_0_.

### Mendelian Randomization

To explore possible causal effects of GGE genome-wide loci on EEG background oscillations, we conducted Mendelian Randomization analyses using GGE and EEGs summary statistics data. 228 SNPs significantly associated with GGE (p<5×10^-8^) were extracted from both the GGE and EEG GWASs. The summary statistics of 228 SNPs were harmonized to ensure the SNP effect direction corresponded with equal effect alleles across GGE and EEG. We used the “TwosampleMR” package ^29^ in R to perform fixed effects inverse variance-weighted (IVW), weighted median, and MR Egger models. We then performed sensitivity analyses, including horizontal pleiotropic effects estimated by the intercept of MR Egger, residual heterogeneity due to pleiotropy estimated by Cochran’s Q test^30^, and leave-one-out analyses (for the IVW-fixed effect model) to evaluate if any single instrumental variable was driving the results. GSMR (generalized summary-data based mendelian randomisation) analyses were performed using the “GSMR” ^31^ package in R. To that end, first the LD matrix of the selected SNPs was calculated using PLINK ^32^ and GCTA ^33^ within 1000 Genomes Phase 3 data ^23^. The minimum number of instrumental variables in the GSMR model was loosened from 10 to 5 as there were only 8 independent (r^2^<0.01, LD window =10Mb) significant loci identified in the GWAS of GGE (and none in the EEG GWAS). We used default options in GSMR with Heterogeneity in Dependent Instruments (HEIDI) testing for instrumental outliers’ detection. At the end, we repeated GSMR with loosened LD prune thresholds (namely r2<0.1, r2<0.15 and r2<0.2) because GSMR takes LD structure into account by adding the LD matrix. The significance threshold was Bonferroni corrected for all these seven mendelian randomisation models (p=0.05/7=0.007).

### Data availability

GWASs summary statistics used for the current analyses are available online: http://enigma.ini.usc.edu/research/download-enigma-gwas-results/; http://www.epigad.org/gwas_ilae2018_16loci.html.

## Results

### Genetic correlations between epilepsy and oscillatory brain activity

In a total study population of 45,446 subjects (n=8,425 from the EEG and n=37,021 from the epilepsy GWASs), we computed genetic correlations between alpha, beta, delta and theta oscillatory brain activity with focal epilepsy and GGE. We found significant correlations between GGE and beta-power (Rg=0.44 ± s.e. of 0.18; p=0.01) and theta-power (Rg=0.25 ± 0.11; p=0.02; **Figure 1**, upper panel, Supplemental table 1). This was further supported by the correlations between beta and theta-power with the GGE subtypes CAE, JAE and JME: all had similarly high correlation coefficients. We found no genetic correlations between focal epilepsy and any of the EEG phenotypes. We then attempted to replicate the genetic correlations using the unpublished Epi25 dataset and found similar genetic correlations (both in sign and effect size) to the discovery analyses (**Figure 1**, lower panel): GGE correlated with beta power (Rg=0.52 ± 0.21, p=0.01), while the genetic correlation between theta power and GGE paralleled the discovery cohort (albeit not reaching significance: Rg=0.16 ± 0.12; p=0.18). All genetic correlation estimates with focal epilepsy were again non-significant. There was no data was available for GGE subtypes.

**Figure 1:**
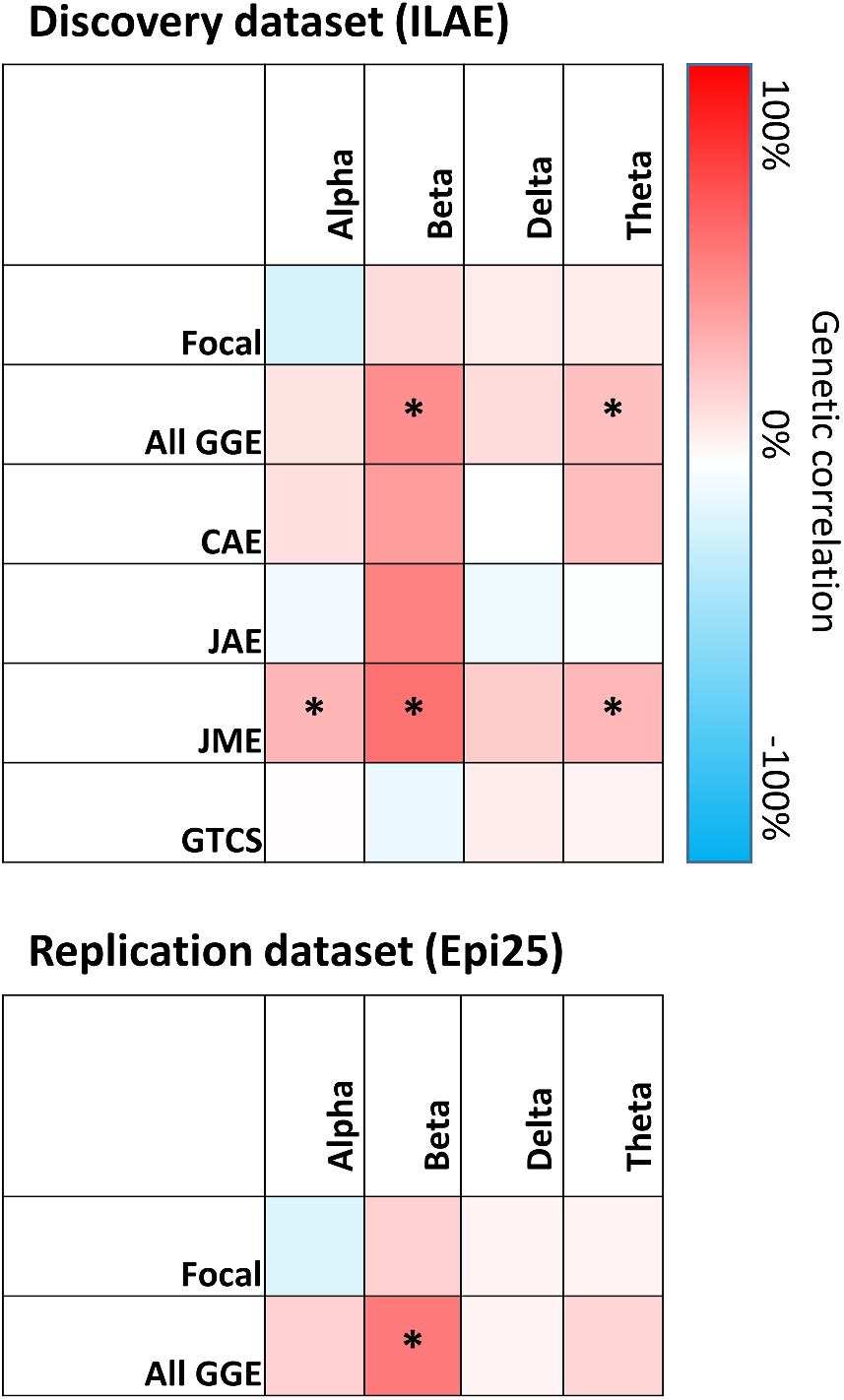
Genetic correlations between EEG frequency bands and epilepsy subtypes. Genetic correlations were calculated by comparing the EEG frequency band GWAS with the ILAE GWAS (upper panel, discovery dataset) and the Epi25 GWAS (lower panel, replication dataset). * p<0.05.

### Oscillatory brain activity polygenic scores are associated with generalized epilepsy

We used polygenic scoring to utilize the full distribution of background EEG associated SNPs to assess whether people with epilepsy have a different polygenic score for specific frequency bands compared to controls. We observed significant positive associations between beta and theta-power PRSs with GGE, in line with the LDSC results (**Figure 2)**. In particular beta power PRSs were strongly associated with GGE (beta=0.11; s.e.= 0.020; p=5.3×10^-8^; explained variance 0.21%; **Figure 2**), which was further supported by significant associations of beta power PRS with its subtypes CAE (beta=0.15; s.e.=0.044; p=8.5×10^-4^) and JME (beta=0.12; s.e.=0.033; p=3.6×10^-4^). Furthermore, of the participants in the GGE case-control cohort, those in the highest 10% decile of beta-power PRS scores were 1.4 fold more likely to have GGE compared to the people in the lowest 10% PRS decile (Supplemental Figure 1; OR: 1.40; 95%CI: 1.18-1.67; p=1.5×10^-4^). When using the independent Epi25 cohort as a replication dataset we found that the directions of effect agreed with the discovery analyses for all associations between EEG PRSs and GGE (p_one-sided_= 0.023; p_two-sided_= 0.046; supplemental Figure 2). EEG PRSs were not significantly different between people with focal epilepsy and controls.

**Figure 2.**
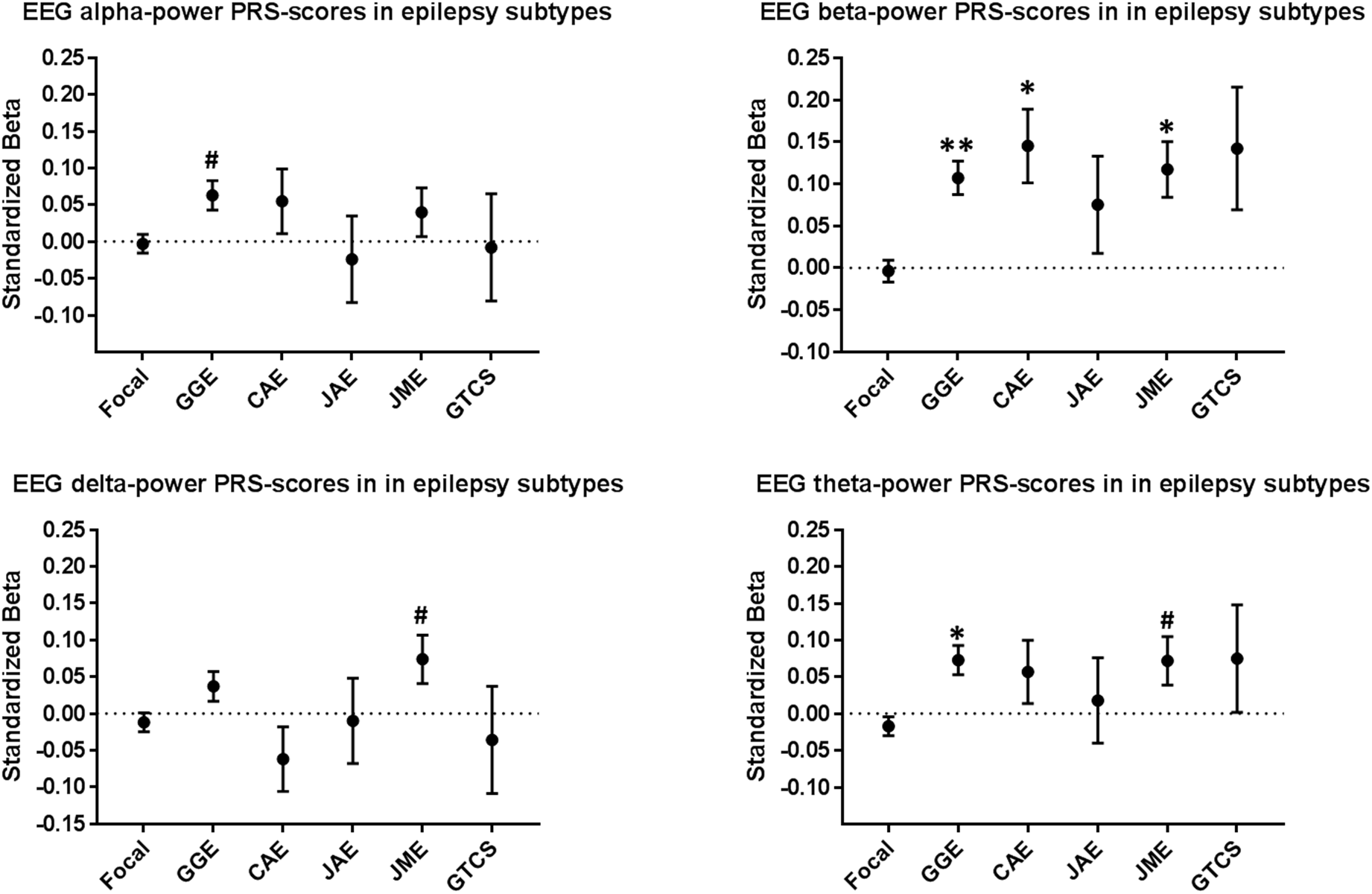
A: Beta and theta power EEG oscillation polygenic risk scores (PRSs) are associated with generalized epilepsy but not with focal epilepsy. The ‘best-fit’ P-value threshold (P_t_) was chosen based on the most significant association with GGE, which was then applied to all other epilepsy subtypes. The number of SNPs included in each model were: 2,670 for Alpha power (P_t_ =0.0105), 10,861 for Beta power (P_t_=0.06245), 8,182 for Delta power (P_t_ =0.0446), and 3,833 for Theta power (P_t_=0.01665). Logistic regression analyses were performed to assess the association between the PRSs and the different epilepsy subtypes, corrected for sex and 10 principal components. #: p<0.05; *: p<0.001; **: P<10^-7^. Focal = focal epilepsy; GGE = genetic generalized epilepsy; CAE (childhood absence epilepsy), JAE (juvenile absence epilepsy), JME (juvenile myoclonic epilepsy) and GTCS (generalized tonic-clonic seizures only) are GGE subtypes.

**Figure 3.**
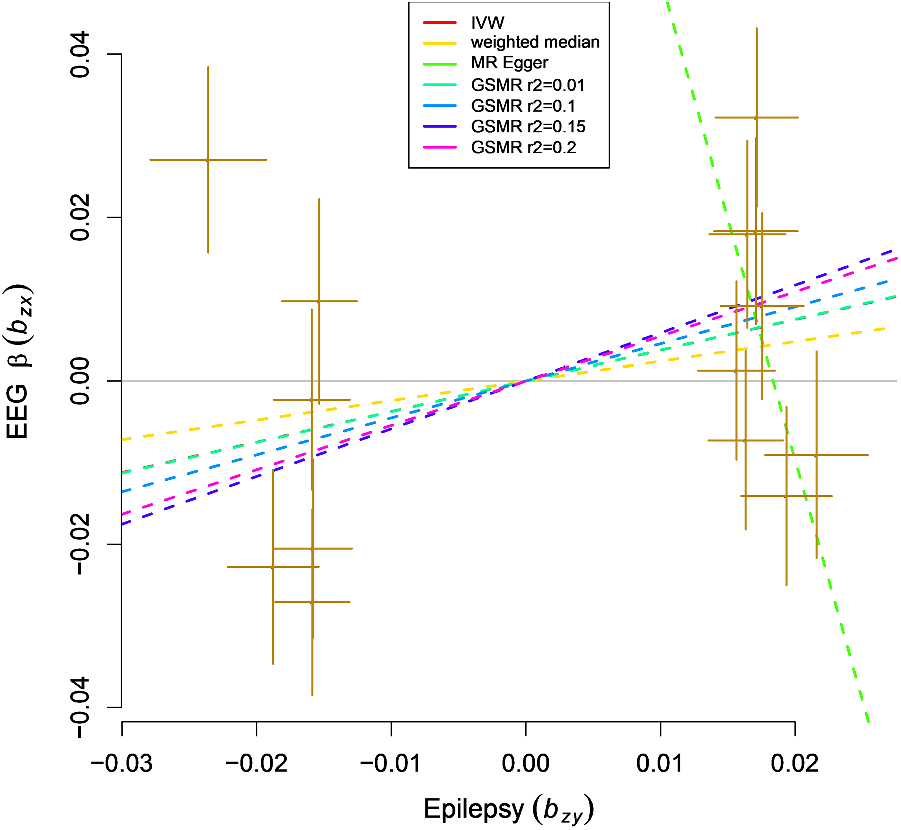
Scatter plot of instrumental SNPs associated with generalized epilepsy tested for EEG beta oscillations. The instrumental variables passed Heterogeneity in Dependent Instruments (HEIDI) testing for instrumental outliers from GSMR detection with multiple LD prune thresholds. Lines in red, yellow, light green and dark green represent β for fixed effect IVW, weighted median, MR Egger, and GSMR models using 8 instruments (r^2^<0.01). The lines in blue purple and pink represent βs from GSMR model with 11 (r^2^<0.1), 12 (r^2^<0.15) and 14 (r^2^<0.2) instruments in GSMR.

### Mendelian Randomization analyses

Mendelian randomisation (MR) analyses were performed to assess potential causative relationships between background EEG and GGE. Eight GGE-associated SNPs were selected as instrumental variables at a strict LD prune threshold (r^2^<0.01, LD window =10Mb). These were used in fixed effects IVW, weighted median, MR Egger, and GSMR (r^2^<0.01) models. After loosening the LD threshold, 11 (r^2^<0.1), 12 (r^2^<0.15) and 14 (r^2^<0.2) SNPs were selected as instrumental variables for GSMR models. Causal effects of GGE loci on beta oscillations were found at the LD r^2^ < 0.15 and <0.2 thresholds (OR=1.79, CI=1.189-2.707, P-value= 5.2×10^-3^; and OR=1.723, CI=1.180-2.516, P-value=4.8×10^-3^, respectively, **Figure 4**, Supplemental Table 4, Supplemental Figure 3). Significant heterogeneity was detected in the fixed-effect IVW model (Q-statistics=18.188, df=7, p= 0.01) and MR Egger model (Q-statistics=14.594, df=6, p= 0.02). No SNPs altered the pooled β coefficient in the leave-one-out sensitivity analysis (β =0.374, p= 0.314) in the fixed-effect IVW model. We found no evidence of horizontal pleiotropic effects. Similarly, the HEIDI test detected no SNPs as pleiotropic outliers.

## Discussion

Here we leveraged the largest currently available GWASs to assess shared genetic underpinnings of epilepsy and of background EEG oscillations. In particular, we found strong genetic relationships between GGE and beta-power osccillations, which were replicated in an independent sample.

Previous studies comparing EEG background oscillations between epilepsy patients and controls are inconsistent: some show increased power in all frequency bands (alpha, beta, delta, theta), whereas others show only increases in specific frequency bands or even decreases in power ^11–16^. This heterogeneity likely reflects multiple variables that are difficult to control for in clinical studies, such as AED usage, sleep deprivation, influence of (inter-) ictal epileptic brain activity, EEG processing and electrode placement. We overcame such limitations by determining the genetic underpinnings of EEG frequency bands in people without epilepsy who are AED-naïve, and with a consistent electrode placement and signal processing. We applied several statistical models to assess this overlap and found that people with generalized, but not focal, epilepsy carry a relative abundance of genetic variation associated with higher beta oscillations. Mendelian Randomisation analyses pointed to causal effects of genetic liability to GGE on beta power.

We did not find genetic correlations between background EEG and focal epilepsy. Although power was limited for this analysis, this finding is consistent with the low contribution of common genetic variants in focal epilepsy and the lack of genetic overlap between focal and generalized epilepsy^10^. Focal epilepsy is likely to represent a more heterogenous group of different causes of epilepsy, many of which do not have a primary genetic cause (e.g. symptomatic epilepsy after traumatic brain injury). Moreover, focal epilepsy by definition only affects one part of the brain, and is therefore less likely to be associated by germline genetic variation and background EEG oscillations which most likely affect the whole brain. In contrast to focal epilepsy, the EEG discharges that characterize generalized epilepsy are dependent on the thalomocortical system 5,17. Similarly, background oscillations have been functionally attributed to the thalamocortical system ^18, 19^, suggesting that thalamocortical functioning could represent a common neurobiological mechanisms reflecting overall brain excitability which influences both GGE risk and (beta-power) background oscillations.

Our results should be interpreted in light of several limitations. First, we are aware of the possible advantages of using Genome Complex Trait Analysis (GCTA) relative to LDSC but since no subject-level genotype data are available for the EEG GWAS, we restricted our genetic correlation estimates LDSC, which is based on summary statistics. LDSC has proven to be a reliable method for genetic correlation estimates and results between LDSC and GCTA have proven consistent. Second, we found that the same genetic variants underlie both GGE and beta-power oscillations, but our study does not prove that people with GGE indeed have altered background oscillations, since we did not have EEG measurements of people with epilepsy in this study. Third, only one-way Mendelian Randomisation analyses were performed due to lack of genome-wide significant loci in the EEG GWAS. We therefore cannot exclude the possibility of bidirectional causality between EEG and GGE. Fourth, we had insufficient data available to carry out subgroup analyses on subjects with non-lesional focal epilepsy patients.

Altogether, our results point to shared biological mechanisms underlying background EEG oscillations and the susceptibility for generalized seizures. Our findings thus open avenues to investigate the clinical utility of background oscillations in genetic generalized epilepsy. Potentially, prospective studies could confirm whether altered beta-oscillatons could be a prodromal state of GGE or whether aberrant beta oscillations constitute a feature of epilepsy. Future studies may also integrate transcranial magnetic stimulation – EEG (TMS-EEG) and/or event-related potentials to examine whether beta and theta powers correlate with altered brain excitability in subjects with high epilepsy liability. Furthermore, machinelearning studies integrating background EEG with other sources of clinical and demographic data may one day increase diagnostic accuracy of epilepsy.

### Key points box

- 1: Genetic correlation studies show shared genetic underpinnings between GGE and power of background oscillations in the beta-frequency band.
- 2: Polygenic risk score analyses show that subjects with more beta-power associated genetic variants have an increased risk of having GGE.
- 3: Mendelian randomization analyses suggest a causal relationship of GGE genetic liability on beta oscillations.

## Acknowledgements

We are grateful to the patients and volunteers who participated in this research. We thank the following clinicians and research scientists for their contribution through sample collection (cases and controls), data analysis, and project support: Geka Ackerhans, Muna Alwaidh, R E Appleton, Willem Frans Arts, Guiliano Avanzini, Paul Boon, Sarah Borror, Kees Braun, Oebele Brouwer, Hans Carpay, Karen Carter, Peter Cleland, Oliver C Cockerell, Paul Cooper, Celia Cramp, Emily de los Reyes, Chris French, Catharine Freyer, William Gallentine, Michel Georges, Peter Goulding, Micheline Gravel, Rhian Gwilliam, Lori Hamiwka, Steven J Howell, Adrian Hughes, Aatif Husain, Monica Islam, Floor Jansen, Mary Karn, Mark Kellett, Ditte B Kjelgaard, Karl Martin Klein, Donna Kring, Annie WC Kung, Mark Lawden, Jo Ellen Lee, Benjamin Legros, Leanne Lehwald, Edouard Louis, Colin HT Lui, Zelko Matkovic, Jennifer McKinney, Brendan McLean, Mohamad Mikati, Bethanie Morgan-Followell, Wim Van Paesschen, Anup Patel, Manuela Pendziwiat, Marcus Reuber, Richard Roberts, Guy Rouleau, Cathy Schumer, B Sharack, Kevin Shianna, NC Sin, Saurabh Sinha, Laurel Slaughter, Sally Steward, Deborah Terry, Chang-Yong Tsao, TH Tsoi, Patrick Tugendhaft, Jaime-Dawn Twanow, Jorge Vidaurre, Sarah Weckhuysen, Pedro Weisleder, Kathleen White, Virginia Wong, Raju Yerra, Jacqueline Yinger and all contributing clinicians from the Department of Clinical and Experimental Epilepsy at the National Hospital for Neurology and Neurosurgery and UCL Institute of Neurology.

We would like to thank the Ming Fund for providing funding for R.S. This work was in part supported by an award by a Translational Research Scholars award from the Health Research Board of Ireland (C.D.W.), by research grants from Science Foundation Ireland (SFI) (16/RC/3948 and 13/CDA/2223) and co-funded under the European Regional Development Fund and by FutureNeuro industry partners. Further funding sources include: Wellcome Trust (grant 084730); Epilepsy Society, UK, NIHR (08-08-SCC); GIHE: NIH R01- NS-49306-01 (R.J.B.); NIH R01-NS-053998 (D.H.L); GSCFE: NIH R01-NS-064154-01 (R.J.B. and Ha.Ha.); NIH: UL1TR001070, Development Fund from The Children’s Hospital of Philadelphia (Ha.Ha.); NHMRC Program Grant ID: 1091593 (S.F.B., I.E.S., K.L.O., and K.E.B.); The Royal Melbourne Hospital Foundation Lottery Grant (S.P.); The RMH Neuroscience Foundation (T.J.O’B.); European Union’s Seventh Framework Programme (FP7/2007-2013) under grant agreement n° 279062 (EpiPGX) and 602102, Department of Health’s NIHR Biomedical Research Centers funding scheme, European Community (EC: FP6 project EPICURE: LSHM-CT2006-037315); German Research Foundation (DFG: SA434/4-1/4-26-1 (Th.Sa.), WE4896/3-1); EuroEPINOMICS Consortium (European Science Foundation/DFG: SA434/5-1, NU50/8-1, LE1030/11-1, HE5415/3-1 (Th.Sa., P.N., H.L., I.H.), RO 3396/2- 1); the German Federal Ministry of Education and Research, National Genome Research Network (NGFNplus/EMINet: 01GS08120, and 01GS08123 (Th.Sa., H.L.); IntenC, TUR 09/I10 (Th.Sa.)); The Netherlands National Epilepsy Fund (grant 04-08); EC (FP7 project EpiPGX 279062). Research Grants Council of the Hong Kong Special Administrative Region, China project numbers HKU7623/08 M (S.S.C, P.K., L.W.B., P.C.S), HKU7747/07 M (S.S.C., P.C.S.) and CUHK4466/06 M (P.K., L.B). Collection of Belgian cases was supported by the Fonds National de la Recherche Scientifique, Fondation Erasme, Université Libre de Bruxelles. GlaxoSmithKline funded the recruitment and data collection for the GenEpA Consortium samples. We acknowledge the support of Nationwide Children’s hospital in Columbus, Ohio, USA. The Wellcome Trust (WT066056) and The NIHR Biomedical Research Centres Scheme (P31753) supported UK contributions. Further support was received through the Intramural Research Program of the Eunice Kennedy Shriver National Institute of Child Health and Human Development (Contract: N01HD33348). The project was also supported by the popgen 2.0 network through a grant from the German Ministry for Education and Research (01EY1103). Parts of the analysis of this work were performed on resources of the High Performance Center of the University of Luxembourg and Elixir-Luxembourg. The KORA study was initiated and financed by the Helmholtz Zentrum München – German Research Center for Environmental Health, which is funded by the German Federal Ministry of Education and Research (BMBF) and by the State of Bavaria. Furthermore, KORA research was supported within the Munich Center of Health Sciences (MC-Health), Ludwig-Maximilians-Universität, as part of LMUinnovativ. The International League Against Epilepsy (ILAE) facilitated the Consortium through the Commission on Genetics and by financial support; however, the opinions expressed in the manuscript do not necessarily represent the policy or position of the ILAE.

## Author contributions statement

RS, JJL, DS and BPCK contributed to the conception and design of the study.

RS, JJL, BDL, DS and BPCK contributed to the acquisition and analysis of data.

RS, JJL, BDL, CL, DL, AS, DS, JAC, KR, RARB, KPJB, FEJ, DS and BPJK contributed to the drafting of the manuscript and preparing the figures. Members of the ILAE consortium on Complex Epilepsies and EPI25 Collaborative contributed clinical and genetic data.

## Ethical Publication Statement

We confirm that we have read the Journal’s position on issues involved in ethical publication and affirm that this report is consistent with those guidelines.

## References

1 Smit, D. J. A., Posthuma, D., Boomsma, D. I. & De Geus, E. J. C. Heritability of background EEG across the power spectrum. Psychophysiology 42, 691–697, doi:10.1111/j.1469-8986.2005.00352.x (2005).

2 Devinsky, O. et al. Epilepsy. Nature Reviews Disease Primers 4, doi:ARTN1802410.1038/nrdp.2018.24 (2018).

3 Thomas, R. H. & Berkovic, S. F. The hidden genetics of epilepsy-a clinically important new paradigm. Nature Reviews Neurology 10, 283–292, doi:10.1038/nrneurol.2014.62 (2014).

4 Steriade, M. Corticothalamic resonance, states of vigilance and mentation. Neuroscience 101, 243–276, doi:10.1016/s0306-4522(00)00353-5 (2000).

5 Gloor, P. Generalized epilepsy with bilateral synchronous spike and wave discharge. New findings concerning its physiological mechanisms. Electroencephalogr Clin Neurophysiol Suppl, 245–249 (1978).

6 Gloor, P. Generalized epilepsy with spike-and-wave discharge: a reinterpretation of its electrographic and clinical manifestations. The 1977 William G. Lennox Lecture, American Epilepsy Society. Epilepsia 20, 571–588 (1979).

7 Gloor, P. & Fariello, R. G. Generalized epilepsy: some of its cellular mechanisms differ from those of focal epilepsy. Trends Neurosci 11, 63–68 (1988).

8 Steriade, M. Sleep, epilepsy and thalamic reticular inhibitory neurons. Trends Neurosci 28, 317–324, doi:10.1016/j.tins.2005.03.007 (2005).

9 Smit, D. J. A. et al. Genome-wide association analysis links multiple psychiatric liability genes to oscillatory brain activity. Human Brain Mapping 39, 4183–4195, doi:10.1002/hbm.24238 (2018).

10 International League Against Epilepsy Consortium on Complex, E. Genome-wide mega-analysis identifies 16 loci and highlights diverse biological mechanisms in the common epilepsies. Nat Commun 9, 5269, doi:10.1038/s41467-018-07524-z (2018).

11 Clemens, B., Szigeti, G. & Barta, Z. EEG frequency profiles of idiopathic generalised epilepsy syndromes. Epilepsy Research 42, 105–115, doi:Doi 10.1016/S0920-1211(00)00167-4 (2000).

12 Clemens, B. Pathological theta oscillations in idiopathic generalised epilepsy. Clinical Neurophysiology 115, 1436–1441, doi:10.1016/j.clinph.2004.01.018 (2004).

13 Clemens, B. et al. Characteristic distribution of interictal brain electrical activity in idiopathic generalized epilepsy. Epilepsia 48, 941–949, doi:10.1111/j.1528-1167.2007.01030.x (2007).

14 Clemens, B. Valproate decreases EEG synchronization In a use-dependent manner in idiopathic generalized epilepsy. Seizure-European Journal of Epilepsy 17, 224–233, doi:10.1016/j.seizure.2007.07.005 (2008).

15 Clemens, B. et al. Valproate treatment normalizes EEG functional connectivity in successfully treated idiopathic generalized epilepsy patients. Epilepsy Research 108, 1896–1903, doi:10.1016/j.eplepsyres.2014.09.032 (2014).

16 Tikka, S. K., Goyal, N., Umesh, S. & Nizamie, S. H. Juvenile myoclonic epilepsy: Clinical characteristics, standard and quantitative electroencephalography analyses. J Pediatr Neurosci 8, 97–103, doi:10.4103/1817-1745.117835 (2013).

17 Steriade, M. & Contreras, D. Spike-wave complexes and fast components of cortically generated seizures. I. Role of neocortex and thalamus. J Neurophysiol 80, 1439–1455, doi:10.1152/jn.1998.80.3.1439 (1998).

18 Steriade, M., Amzica, F. & Contreras, D. Synchronization of fast (30-40 Hz) spontaneous cortical rhythms during brain activation. J Neurosci 16, 392–417 (1996).

19 Steriade, M., Contreras, D., Amzica, F. & Timofeev, I. Synchronization of fast (30-40 Hz) spontaneous oscillations in intrathalamic and thalamocortical networks. J Neurosci 16, 2788–2808 (1996).

20 Gray, C. M. & Singer, W. Stimulus-specific neuronal oscillations in orientation columns of cat visual cortex. Proc Natl Acad Sci U S A 86, 1698–1702, doi:10.1073/pnas.86.5.1698 (1989).

21 MacKay, W. A. & Mendonca, A. J. Field potential oscillatory bursts in parietal cortex before and during reach. Brain Res 704, 167–174, doi:10.1016/0006-8993(95)01109-9 (1995).

22 Bulik-Sullivan, B. K. et al. LD Score regression distinguishes confounding from polygenicity in genome-wide association studies. Nat Genet 47, 291–295, doi:10.1038/ng.3211 (2015).

23 Genomes Project, C. et al. A global reference for human genetic variation. Nature 526, 68–74, doi:10.1038/nature15393 (2015).

24 Manichaikul, A. et al. Robust relationship inference in genome-wide association studies. Bioinformatics 26, 2867–2873, doi:10.1093/bioinformatics/btq559 (2010).

25 Euesden, J., Lewis, C. M. & O’Reilly, P. F. PRSice: Polygenic Risk Score software. Bioinformatics 31, 1466–1468, doi:10.1093/bioinformatics/btu848 (2015).

26 Demontis, D. et al. Discovery of the first genome-wide significant risk loci for attention deficit/hyperactivity disorder. Nat Genet 51, 63–75, doi:10.1038/s41588-018-0269-7 (2019).

27 Barbeira, A. N. et al. Exploring the phenotypic consequences of tissue specific gene expression variation inferred from GWAS summary statistics. Nat Commun 9, 1825, doi:10.1038/s41467-018-03621-1 (2018).

28 Gamazon, E. R. et al. A gene-based association method for mapping traits using reference transcriptome data. Nat Genet 47, 1091–1098, doi:10.1038/ng.3367 (2015).

29 Hemani, G. et al. The MR-Base platform supports systematic causal inference across the human phenome. Elife 7, doi:10.7554/eLife.34408 (2018).

30 Burgess, S. et al. Using published data in Mendelian randomization: a blueprint for efficient identification of causal risk factors. European Journal of Epidemiology 30, 543–552, doi:10.1007/s10654-015-0011-z (2015).

31 Zhu, Z. H. et al. Causal associations between risk factors and common diseases inferred from GWAS summary data. Nature Communications 9, doi:ARTN 224 10.1038/s41467-017-02317-2 (2018).

32 Purcell, S. et al. PLINK: A tool set for whole-genome association and population-based linkage analyses. American Journal of Human Genetics 81, 559–575, doi:10.1086/519795 (2007).

33 Yang, J., Lee, S. H., Goddard, M. E. & Visscher, P. M. GCTA: a tool for genome-wide complex trait analysis. Am J Hum Genet 88, 76–82, doi:10.1016/j.ajhg.2010.11.011 (2011).

