## supplements for "Shared genetic underpinnings between genetic generalized epilepsy and background EEG oscillations"

**Supporting information - Converging evidence for shared genetic underpinnings of genetic generalized epilepsy and EEG beta power**

Remi Stevelink, MD, MSc ^1,2,*^, Jurjen J. Luykx MD, PhD^3,4,5,*,#^, Bochao D. Lin, PhD^3,4,^ Costin Leu, PhD^6^, Dennis Lal, PhD^6^, Alexander Smith, MA^6^, Dick Schijven, PhD^3,4^, Johannes A. Carpay, MD, PhD^7^, Koen Rademaker, BSc^3^, Roiza A. Rodrigues Baldez, MD^8^, Orrin Devinsky^9^, MD, Kees P.J. Braun MD, PhD^2^, Floor E. Jansen, MD, PhD^2^, ILAE Consortium on Complex Epilepsies, Epi25 Collaborative, Dirk Smit PhD^10,@^ & Bobby P.C. Koeleman, PhD^1,@^

^1^Department of Genetics, UMC Utrecht Brain Center, University Medical Center Utrecht, Utrecht University, Utrecht, the Netherlands

^2^Department of Neurology, UMC Utrecht Brain Center, University Medical Center Utrecht, Utrecht University, Utrecht, the Netherlands

^3^Department of Psychiatry, UMC Utrecht Brain Center, University Medical Center Utrecht, Utrecht University, Utrecht, the Netherlands;

^4^Department of Translational Neuroscience, UMC Utrecht Brain Center, University Medical Center Utrecht, Utrecht University, Utrecht, The Netherlands

^5^GGNet Mental Health, Apeldoorn, the Netherlands

^6^Broad Institute of Harvard and MIT, Cambridge, Massachussets, USA.

^7^Department of Neurology, Tergooi Hospital, Hilversum, the Netherlands

^8^ Clinical Research Laboratory on Neuroinfectious Diseases, Evandro Chagas Clinical Research Institute (IPEC), Fundação Oswaldo Cruz (FIOCRUZ), Rio de Janeiro, RJ, Brazil

^9^ Comprehensive Epilepsy Center, New York University School of Medicine, New York, NY 10016, USA.
^10^ Psychiatry department, Amsterdam Neuroscience, Amsterdam University Medical Center, location AMC, The Netherlands.

^*^ These authors contributed equally to this work;

^@^ These authors jointly directed this work.

^#^ To whom correspondence should be addressed at:

Jurjen Luykx, Department of Psychiatry, Brain Center Rudolf Magnus, University Medical Center Utrecht, Utrecht University, Utrecht, the Netherlands;

**Supporting materials**

**Supplementary Table 1**

**Number of SNPs tested per LDSC genetic correlation analysis.**

| Epilepsy phenotype | Alpha | Beta | Delta | Theta |
| --- | --- | --- | --- | --- |
| ILAE focal | 842636 | 843077 | 842616 | 842616 |
| ILAE GGE | 842636 | 843077 | 842616 | 842616 |
| ILAE CAE | 864641 | 863614 | 863614 | 863614 |
| ILAE JAE | 864641 | 863614 | 863614 | 863614 |
| ILAE JME | 864641 | 863614 | 863614 | 863614 |
| ILAE GTCS | 864641 | 863614 | 863614 | 863614 |
| Epi25 focal | 288911 | 288902 | 288902 | 288902 |
| Epi25 GGE | 288911 | 288902 | 288902 | 288902 |

**Supplementary Table 2**

*How all EEG power spectra are correlated may be found in this correlation matrix of Spearman correlations (N=8,425). All p-values were well below e-17.*

|  | Delta | Theta | Alpha | Beta |
| --- | --- | --- | --- | --- |
| Delta | 1 |  |  |  |
| Theta | 0.7002 | 1 |  |  |
| Alpha | 0.3004 | 0.5422 | 1 |  |
| Beta | 0.4232 | 0.5247 | 0.6252 | 1 |

**Supplemental Table 3**

*Summary table of LDSC genetic correlation analyses (p1=phenotype 1; p2=phenotype 2; rg=genetic correlation; se=standard error; z=z-score; p=p-value; h2_obs, h2_obs_se=observed scale h2 for phenotype 2 and standard error; h2_int, h2_int_se=single-trait LD Score regression intercept for phenotype 2 and standard error; gcov_int_se=cross-trait LD Score regression intercept and standard error. Note: Focal epilepsy showed low SNP heritability and z-score<2, which makes the genetic correlation estimates less reliable.*

| p1 | p2 | rg | se | z | p | h2_obs | h2_obs_se | h2_int | h2_int_se | gcov_int | gcov_int_se |
| --- | --- | --- | --- | --- | --- | --- | --- | --- | --- | --- | --- |
| alpha | All epilepsy | 0.0591 | 0.1286 | 0.4595 | 0.6459 | 0.1327 | 0.0202 | 1.1693 | 0.0125 | 0.0032 | 0.007 |
| alpha | *Focal epilepsy* | *-0.1652* | *0.2784* | *-0.5934* | *0.5529* | *0.0356* | *0.0206* | *1.1704* | *0.0107* | *0.0008* | *0.0069* |
| alpha | GGE | 0.1569 | 0.107 | 1.4664 | 0.1426 | 0.323 | 0.033 | 1.1036 | 0.0134 | 0.005 | 0.0078 |
| alpha | CAE | 0.1274 | 0.1731 | 0.7361 | 0.4617 | 1.005 | 0.22 | 1.0724 | 0.0107 | 0.0106 | 0.007 |
| alpha | JAE | -0.041 | 0.22 | -0.1863 | 0.8522 | 1.1673 | 0.3377 | 1.0486 | 0.0094 | 0.0096 | 0.0074 |
| alpha | JME | 0.2879 | 0.1414 | 2.0358 | 0.0418 | 1.0461 | 0.1711 | 1.0551 | 0.0118 | -0.0031 | 0.0074 |
| alpha | GTCS | 0.0085 | 0.217 | 0.0391 | 0.9688 | 1.8076 | 0.5777 | 1.0297 | 0.0096 | 0.002 | 0.0064 |
| beta | All epilepsy | 0.3032 | 0.1895 | 1.5996 | 0.1097 | 0.1326 | 0.0202 | 1.1694 | 0.0125 | -0.0056 | 0.0075 |
| beta | *Focal epilepsy* | *0.1414* | *0.3661* | *0.3861* | *0.6994* | *0.0356* | *0.0206* | *1.1704* | *0.0107* | *-0.0052* | *0.0069* |
| beta | GGE | 0.4436 | 0.1766 | 2.5124 | **0.012** | 0.3229 | 0.033 | 1.1036 | 0.0134 | -0.004 | 0.0075 |
| beta | CAE | 0.3906 | 0.2376 | 1.6439 | 0.1002 | 1.0047 | 0.2197 | 1.0724 | 0.0107 | 0.0043 | 0.0065 |
| beta | JAE | 0.4928 | 0.304 | 1.6212 | 0.105 | 1.1689 | 0.3375 | 1.0485 | 0.0094 | -0.0038 | 0.0069 |
| beta | JME | 0.5568 | 0.2197 | 2.5341 | **0.0113** | 1.0463 | 0.1711 | 1.055 | 0.0118 | -0.0085 | 0.0071 |
| beta | GTCS | -0.0771 | 0.3153 | -0.2445 | 0.8068 | 1.8064 | 0.5784 | 1.0297 | 0.0096 | 0.0117 | 0.0067 |
| delta | All epilepsy | 0.1162 | 0.1604 | 0.7245 | 0.4688 | 0.1326 | 0.0202 | 1.1694 | 0.0125 | -0.009 | 0.0072 |
| delta | *Focal epilepsy* | *0.0798* | *0.3192* | *0.2498* | *0.8027* | *0.0356* | *0.0206* | *1.1704* | *0.0107* | *-0.0084* | *0.0071* |
| delta | GGE | 0.1392 | 0.1237 | 1.1256 | 0.2603 | 0.3229 | 0.033 | 1.1036 | 0.0134 | -0.0069 | 0.0074 |
| delta | CAE | 0.0007 | 0.1829 | 0.0041 | 0.9967 | 1.0047 | 0.2197 | 1.0724 | 0.0107 | -0.0019 | 0.0066 |
| delta | JAE | -0.0654 | 0.2071 | -0.3158 | 0.7521 | 1.1689 | 0.3375 | 1.0485 | 0.0094 | -0.0003 | 0.0068 |
| delta | JME | 0.2014 | 0.1579 | 1.2756 | 0.2021 | 1.0463 | 0.1711 | 1.055 | 0.0118 | -0.0071 | 0.0073 |
| delta | GTCS | 0.0742 | 0.2609 | 0.2843 | 0.7762 | 1.8064 | 0.5784 | 1.0297 | 0.0096 | -0.0055 | 0.0066 |
| theta | All epilepsy | 0.2269 | 0.1271 | 1.7848 | 0.0743 | 0.1326 | 0.0202 | 1.1694 | 0.0125 | -0.0099 | 0.007 |
| theta | *Focal epilepsy* | *0.0757* | *0.2926* | *0.2587* | *0.7958* | *0.0356* | *0.0206* | *1.1704* | *0.0107* | *-0.0073* | *0.0073* |
| theta | GGE | 0.2498 | 0.1058 | 2.36 | 0.0183 | 0.3229 | 0.033 | 1.1036 | 0.0134 | -0.0034 | 0.007 |
| theta | CAE | 0.2615 | 0.1874 | 1.3952 | 0.163 | 1.0047 | 0.2197 | 1.0724 | 0.0107 | -0.002 | 0.007 |
| theta | JAE | -0.0099 | 0.1943 | -0.0508 | 0.9595 | 1.1689 | 0.3375 | 1.0485 | 0.0094 | 0.0056 | 0.007 |
| theta | JME | 0.2838 | 0.1344 | 2.112 | 0.0347 | 1.0463 | 0.1711 | 1.055 | 0.0118 | -0.0082 | 0.0068 |
| theta | GTCS | 0.0438 | 0.2376 | 0.1842 | 0.8539 | 1.8064 | 0.5784 | 1.0297 | 0.0096 | 0.0017 | 0.0067 |

**Supplemental Table 4.**

Mendelian Randomization results (in bold: significant results after multiple testing correction).

| Outcome | Model | N instrument | N* | OR (95%CI) | P |
| --- | --- | --- | --- | --- | --- |
| Alpha | IVW-fixed | 8 | 5 | 1.470(0.935-2.312) | 0.095 |
|  | Weighted median | 8 | 5 | 2.017(1.069-3.808) | 0.031 |
|  | MR egger | 8 | 5 | 0 (0-182.998) | 0.546 |
|  | MR egger (intercept) | 8 | 5 | 0.053 ± 0.068 | 0.487 |
|  | GSMR (r2<0.01) | 8 | 5 | 1.473 (0.927-2.339) | 0.101 |
|  | GSMR (r2<0.1) | 11 | 7 | 1.173(0.820-1.678) | 0.381 |
|  | GSMR (r2<0.15) | 12 | 8 | 1.238 (0.874-1.755) | 0.228 |
|  | GSMR (r2<0.2) | 14 | 9 | 1.346 (0.953-1.900) | 0.091 |
| Beta | IVW-fixed | 8 | 5 | 1.454 (0.925-2.286) | 0.105 |
|  | Weighted median | 8 | 5 | 1.270 (0.624-2.583) | 0.509 |
|  | MR egger | 8 | 5 | 0.003 (0-69.362) | 0.297 |
|  | MR egger (intercept) | 8 | 5 | 0.108 ± 0.089 | 0.269 |
|  | GSMR (r2<0.01) | 8 | 5 | 1.456 (0.920-2.303) | 0.108 |
|  | GSMR (r2<0.1) | 11 | 6 | 1.572 (1.049-2.355) | 0.028 |
|  | **GSMR (r2<0.15)** | **12** | **8** | **1.79 (1.189-2.707)** | **0.0052** |
|  | **GSMR (r2<0.2)** | **14** | **9** | **1.723 (1.180-2.516)** | **0.0048** |
| Delta | IVW-fixed | 8 | 5 | 1.387 (0.88292.181) | 0.156 |
|  | Weighted median | 8 | 5 | 1.064 (0.551-2.057) | 0.852 |
|  | MR egger | 8 | 5 | 0.035 (0-225.458) | 0.481 |
|  | MR egger (intercept) | 8 | 5 | 0.064± 0.077 | 0.441 |
|  | GSMR (r2<0.01) | 8 | 5 | 1.387 (0.882-2.181) | 0.156 |
|  | GSMR (r2<0.1) | 11 | 7 | 1.295 (0.906-1.849) | 0.154 |
|  | GSMR (r2<0.15) | 12 | 8 | 1.360 (0.958-1.931) | 0.085 |
|  | GSMR (r2<0.2) | 14 | 8 | 1.360 (0.958-1.932) | 0.085 |
| Theta | IVW-fixed | 8 | 6 | 1.322 (0.841-2.078) | 0.227 |
|  | Weighted median | 8 | 6 | 1.242 (0.707-2.181) | 0.450 |
|  | MR egger | 8 | 6 | 0.038 (0-25.341) | 0.363 |
|  | MR egger (intercept) | 8 | 6 | 0.061±0.057 | 0.324 |
|  | GSMR (r2<0.01) | 8 | 6 | 1.322 (0.839-2.082) | 0.228 |
|  | GSMR (r2<0.1) | 11 | 9 | 1.313 (0.917-1.881) | 0.137 |
|  | GSMR (r2<0.15) | 12 | 10 | 1.434 (1.006-2.045) | 0.046 |
|  | GSMR (r2<0.2) | 14 | 10 | 1.431 (1.010-2.027) | 0.044 |

Note: N* is the number of instrumental SNPs have same direction of effects on both exposure (epilepsy) and outcome (EEG).

**Supplemental Figure 1**

*Polygenic risk score (PRS) analyses show that higher beta-power PRS is associated with an increased likelihood of having GGE. All subjects were divided into 10 deciles based on their beta-power PRS scores. Logistic regression analyses were performed to quantify the increased risk of having GGE between every decile compared to the lowest decile (0-10%) as a reference. The odds ratios of these analyses are displayed on the Y-axis. #: P<0.05; *: P<0.001.*

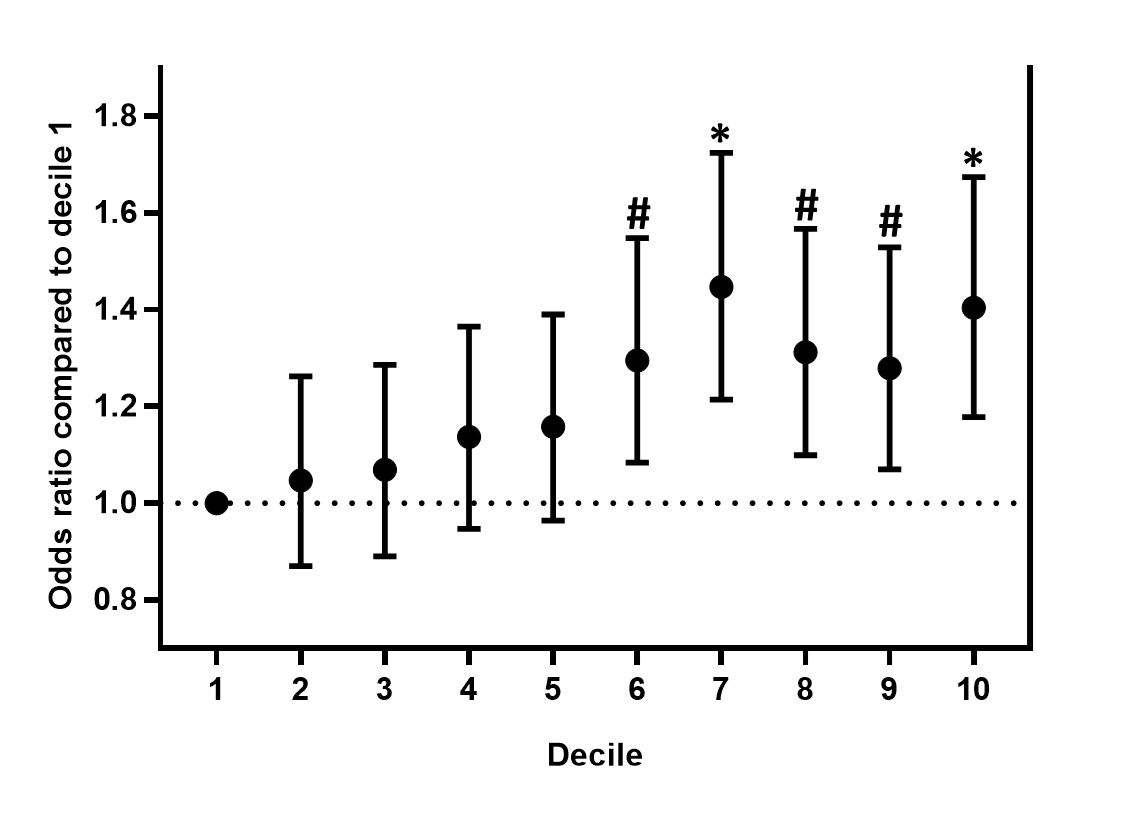

**Supplemental Figure 2**

*Polygenic risk score (PRS) analyses results in the Epi25 replication cohort. All directions of effect agreed with the PRS analyses in the discovery cohort. Theta-power PRSs were most strongly associated with GGE in this replication cohort (p=0.008).*

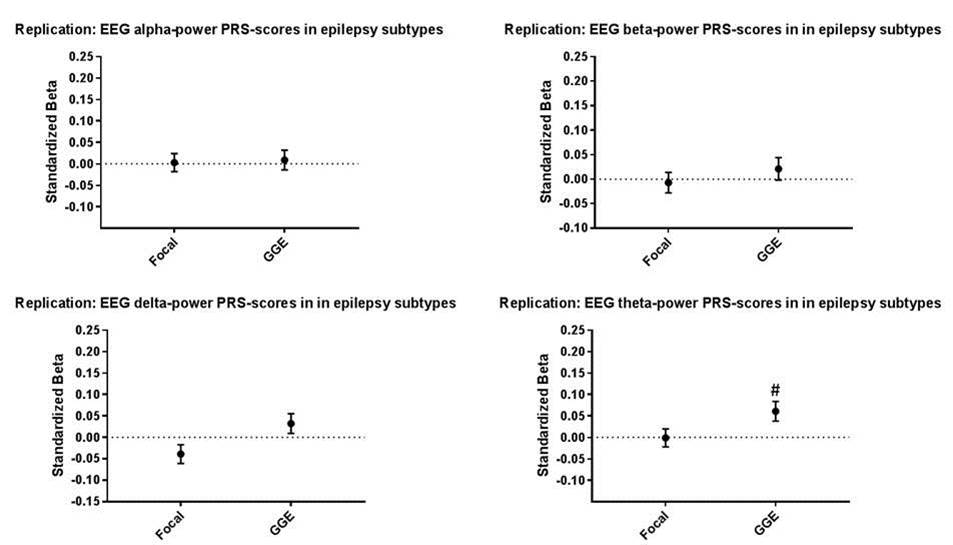

**Supplemental Figure 3**

*Scatter plot of instrumental SNPs associated with generalized epilepsy against EEG alpha, delta and theta oscillations. The instrumental variables passed the Heterogeneity in Dependent Instruments (HEIDI) testing for instrumental outliers from GSMR detection with multiple LD prune thresholds. Lines in red, yellow, light green and dark green represent β for fixed effect IVW, weighted median, MR Egger, and GSMR models using 8 instruments (r2<0.01). The lines in blue purple and pink represent βs from GSMR model with 11 (r2<0.1), 12 (r2<0.15)and 14 (r2<0.2) instruments in GSMR.*
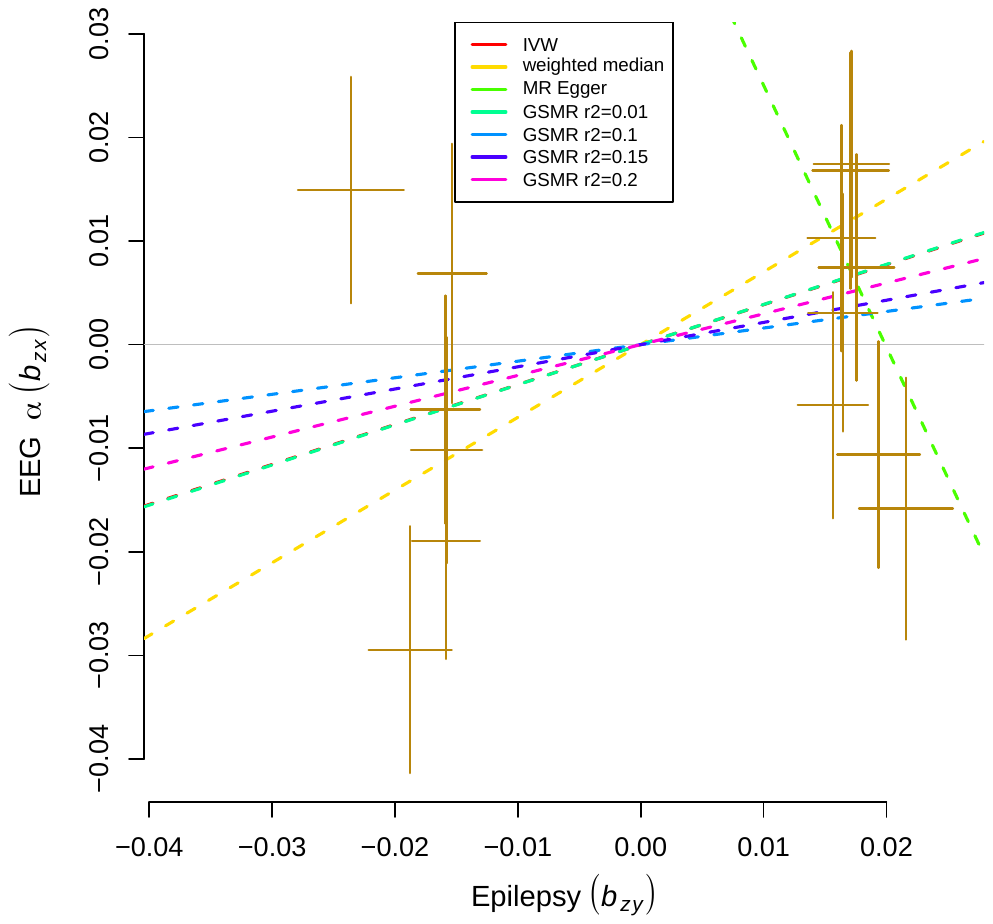

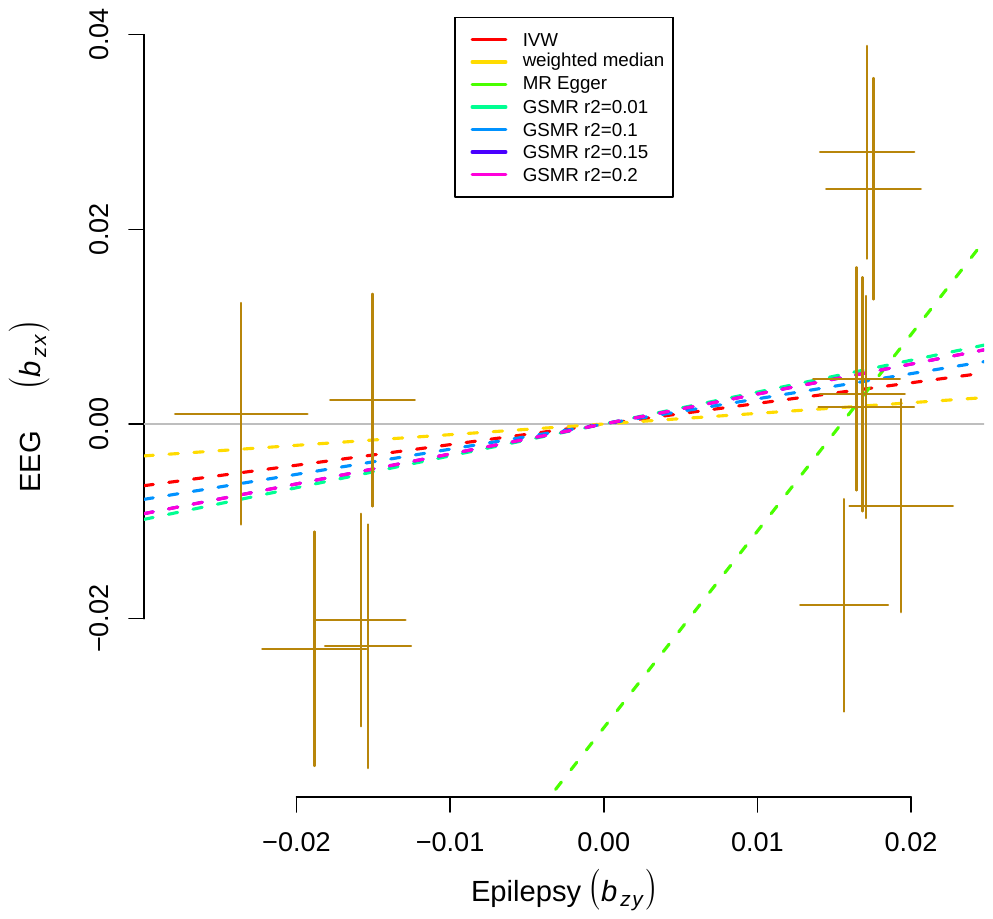

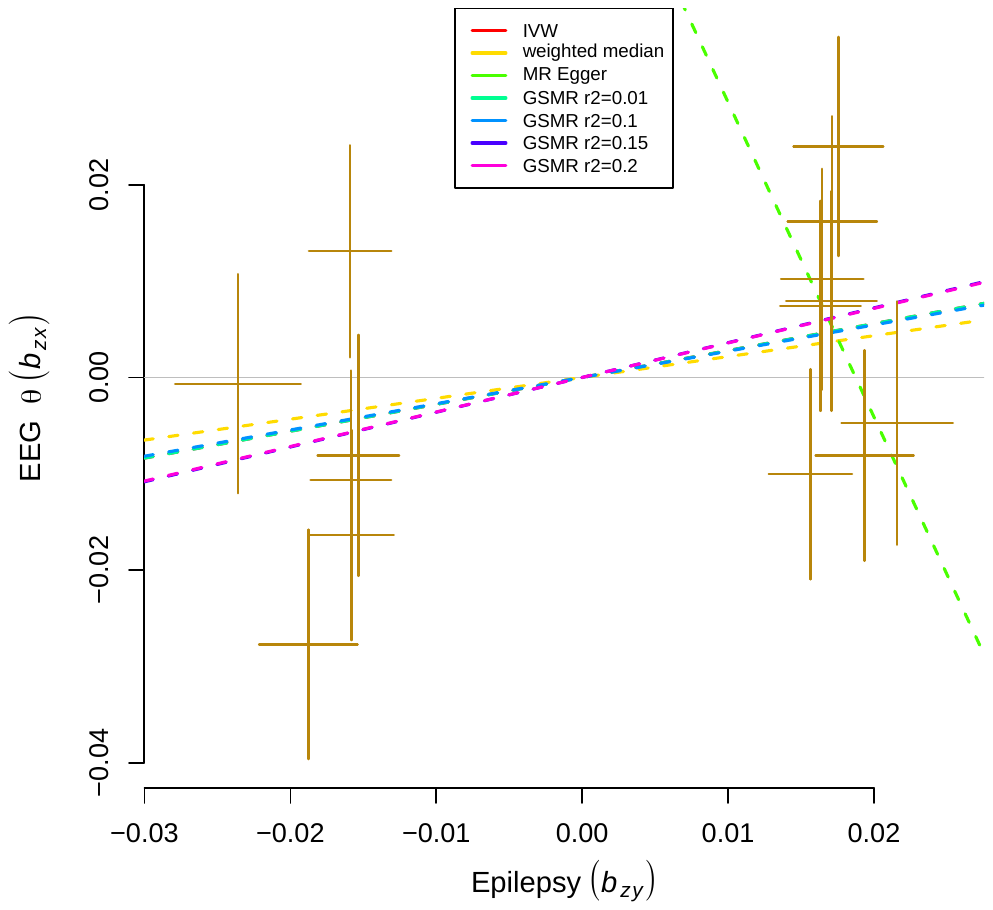

**Appendix**: ILAE consortium author names:

The International League Against Epilepsy Consortium on Complex Epilepsies

*Members listed in alphabetical order:*

Bassel Abou-Khalil^1^, Pauls Auce^2, 3^, Andreja Avbersek^4^, Melanie Bahlo^5-7^, David J Balding^8, 9^, Thomas Bast^10, 11^, Larry Baum^12-14^, Albert J Becker^15^, Felicitas Becker^16, 17^ Bianca Berghuis^18^, Samuel F Berkovic^19^, Katja E Boysen^19^, Jonathan P Bradfield^20, 21^, Lawrence C Brody^22^, Russell J Buono^20, 23, 24^, Ellen Campbell^25^, Gregory D Cascino^26^, Claudia B Catarino^4^, Gianpiero L Cavalleri^27, 28^, Stacey S Cherny^13, 29^, Krishna Chinthapalli^4^, Alison J Coffey^30^, Alastair Compston^31^, Antonietta Coppola^32, 33^, Patrick Cossette^34^, John J Craig^35^, Gerrit-Jan de Haan^36^, Peter De Jonghe^37, 38^, Carolien G F de Kovel^39^, Norman Delanty^27, 28, 40^, Chantal Depondt^41^, Orrin Devinsky^42^, Dennis J Dlugos^43^, Colin P Doherty^28, 44^, Christian E Elger^45^, Johan G Eriksson^46^, Thomas N Ferraro^23, 47^, Martha Feucht^48^, Ben Francis^49^, Andre Franke^50^, Jacqueline A French^51^, Saskia Freytag^5^, Verena Gaus^52^, Eric B Geller^53^, Christian Gieger^54, 55^, Tracy Glauser^56^, Simon Glynn^57^, David B Goldstein^58, 59^, Hongsheng Gui^13^, Youling Guo^13^, Kevin F Haas^1^, Hakon Hakonarson^20, 60^, Kerstin Hallmann^45, 61^, Sheryl Haut^62^, Erin L Heinzen^58, 59^, Ingo Helbig^43, 63^, Christian Hengsbach^16^, Helle Hjalgrim^64, 65^, Michele Iacomino^33^, Andrés Ingason^66^, Jennifer Jamnadas-Khoda^4, 67^, Michael R Johnson^68^, Reetta Kälviäinen^69, 70^, Anne-Mari Kantanen^69^, Dalia Kasperavičiūte^4^, Dorothee Kasteleijn-Nolst Trenite^39^, Heidi E Kirsch^71^, Robert C Knowlton^72^, Bobby P C Koeleman^39^, Roland Krause^73^, Martin Krenn^74^, Wolfram S Kunz^45^, Ruben Kuzniecky^75^, Patrick Kwan^12, 76, 77^, Dennis Lal^78^, Yu-Lung Lau^79^, Anna-Elina Lehesjoki^80^, Holger Lerche^16^, Costin Leu^4, 78, 81^, Wolfgang Lieb^82^, Dick Lindhout^36, 39^, Warren D Lo^83^, Iscia Lopes-Cendes^84, 85^, Daniel H Lowenstein^71^, Alberto Malovini^86^, Anthony G Marson^2^, Thomas Mayer^87^, Mark McCormack^27^, James L Mills^88^, Nasir Mirza^2^, Martina Moerzinger^48^, Rikke S Møller^64, 65, 89^, Anne M Molloy^90^, Hiltrud Muhle^63^, Mark Newton^91^, Ping-Wing Ng^92^, Markus M Nöthen^93^, Peter Nürnberg^94^, Terence J O’Brien^76, 77^, Karen L Oliver^19^, Aarno Palotie^95, 96^, Faith Pangilinan^22^, Sarah Peter^73^, Slavé Petrovski^76, 97^, Annapurna Poduri^98^, Michael Privitera^99^, Rodney Radtke^100^, Sarah Rau^16^, Philipp S Reif^101, 102^, Eva M Reinthaler^74^, Felix Rosenow^101, 102^, Josemir W Sander^4, 36, 103^, Thomas Sander^52, 94^, Theresa Scattergood^104^, Steven C Schachter^105^, Christoph J Schankin^106^, Ingrid E Scheffer^19, 107^, Bettina Schmitz^52^, Susanne Schoch^15^, Pak C Sham^13^, Jerry J Shih^108^, Graeme J Sills^2^, Sanjay M Sisodiya^4, 103^, Lisa Slattery^109^, Alexander Smith^78^, David F Smith^3^, Michael C Smith^110^, Philip E Smith^111^, Anja C M Sonsma^39^, Doug Speed^8, 112^, Michael R Sperling^113^, Bernhard J Steinhoff^10^, Ulrich Stephani^63^, Remi Stevelink^39^, Konstantin Strauch^114, 115^, Pasquale Striano^116^, Hans Stroink^117^, Rainer Surges^45^, K Meng Tan^76^, Liu Lin Thio^118^, G Neil Thomas^119^, Marian Todaro^76^, Rossana Tozzi^120^, Maria S Vari^116^, Eileen P G Vining^121^, Frank Visscher^122^, Sarah von Spiczak^63^, Nicole M Walley^58, 123^, Yvonne G Weber^16^, Zhi Wei^124^, Ruta Mameniskiene^125^, Judith Weisenberg^118^, Christopher D Whelan^27^, Peter Widdess-Walsh^53^, Markus Wolff^125^, Stefan Wolking^16^, Wanling Yang^79^, Federico Zara^33^, Fritz Zimprich^74^

1. Vanderbilt University Medical Center, Nashville, TN 37232, USA.

2. Department of Molecular and Clinical Pharmacology, University of Liverpool, Liverpool L69 3GL, UK.

3. The Walton Centre NHS Foundation Trust, Liverpool L9 7LJ, UK.

4. Department of Clinical and Experimental Epilepsy, UCL Institute of Neurology, Queen Square, London WC1N 3BG, UK.

5. Population Health and Immunity Divison, The Walter and Eliza Hall Institute of Medical Research, Parkville 3052, Australia.

6. Department of Biology, University of Melbourne, Parkville 3010, Australia.

7. School of Mathematics and Statistics, University of Melbourne, Parkville 3010, Australia.

8. UCL Genetics Institute, University College London, London WC1E 6BT, UK.

9. Melbourne Integrative Genomics, University of Melbourne, Parkville 3052, Australia.

10. Epilepsy Center Kork, Kehl-Kork 77694, Germany.

11. Medical Faculty of the University of Freiburg, Freiburg 79085, Germany.

12. Centre for Genomic Sciences, The University of Hong Kong, Hong Kong.

13. Department of Psychiatry, The University of Hong Kong, Hong Kong.

14. The State Key Laboratory of Brain and Cognitive Sciences, University of Hong Kong, Hong Kong, China.

15. Section for Translational Epilepsy Research, Department of Neuropathology, University of Bonn Medical Center, Bonn 53105, Germany.

16. Department of Neurology and Epileptology, Hertie Institute for Clinical Brain Research, University of Tübingen, Tübingen 72076, Germany.

17. Department of Neurology, University of Ulm, Ulm 89081, Germany.

18. Stichting Epilepsie Instellingen Nederland (SEIN), Zwolle 8025 BV, The Netherlands.

19. Epilepsy Research Centre, University of Melbourne, Austin Health, Heidelberg 3084, Australia.

20. Center for Applied Genomics, The Children's Hospital of Philadelphia, Philadelphia, PA 19104, USA.

21. Quantinuum Research LLC, San Diego, CA 92101, USA.

22. National Human Genome Research Institute, National Institutes of Health, Bethesda, MD 20892, USA.

23. Department of Biomedical Sciences, Cooper Medical School of Rowan University Camden, NJ 08103, USA.

24. Department of Neurology, Thomas Jefferson University Hospital, Philadelphia, PA 19107, USA.

25. Belfast Health and Social Care Trust, Belfast BT9 7AB, UK.

26. Division of Epilepsy, Department of Neurology, Mayo Clinic, Rochester, MN 55902, USA.

27. Department of Molecular and Cellular Therapeutics, The Royal College of Surgeons in Ireland, Dublin 2, Ireland.

28. The FutureNeuro Research Centre, Dublin 2, Ireland.

29. Department of Epidemiology and Preventive Medicine, School of Public Health, Sackler Faculty of Medicine, Tel Aviv University, Tel Aviv 6997801, Israel.

30. The Wellcome Trust Sanger Institute, Hinxton, Cambridge CB10 1SA, UK.

31. Department of Clinical Neurosciences, Cambridge Biomedical Campus, Cambridge CB2 0SL, UK.

32. Department of Neuroscience, Reproductive and Odontostomatological Sciences, University Federico II, Naples 80138, Italy.

33. Laboratory of Neurogenetics and Neurosciences, Institute G. Gaslini, Genova 16148, Italy.

34. Department of Neurosciences, Université de Montréal, Montréal, CA 26758, Canada.

35. Department of Neurology, Royal Victoria Hospital, Belfast Health and Social Care Trust, Grosvenor Road, Belfast BT12 6BA, UK.

36. Stichting Epilepsie Instellingen Nederland (SEIN), Heemstede 2103 SW, The Netherlands.

37. Neurogenetics Group, Center for Molecular Neurology, VIB and Laboratory of Neurogenetics, Institute Born-Bunge, University of Antwerp, Antwerp 2610, Belgium.

38. Department of Neurology, Antwerp University Hospital, Edegem 2650, Belgium.

39. Department of Genetics, University Medical Center Utrecht, Utrecht 3584 CX, The Netherlands.

40. Division of Neurology, Beaumont Hospital, Dublin D09 FT51, Ireland.

41. Department of Neurology, Hôpital Erasme, Université Libre de Bruxelles, Bruxelles 1070, Belgium.

42. Comprehensive Epilepsy Center, New York University School of Medicine, New York, NY 10016, USA.

43. Department of Neurology, The Children's Hospital of Philadelphia, Philadelphia, PA 19104, USA.

44. Neurology Department, St. James’s Hospital, Dublin D03 VX82, Ireland.

45. Department of Epileptology, University of Bonn Medical Centre, Bonn 53127, Germany.

46. Department of General Practice and Primary Health Care, University of Helsinki and Helsinki University Hospital, Helsinki 0014, Finland.

47. Department of Pharmacology and Psychiatry, University of Pennsylvania Perlman School of Medicine, Philadelphia, PA 19104, USA.

48. Department of Pediatrics and Neonatology, Medical University of Vienna, Vienna 1090, Austria.

49. Department of Biostatistics, University of Liverpool, Liverpool L69 3GL, UK.

50. Institute of Clinical Molecular Biology, Christian-Albrechts-University of Kiel, University Hospital Schleswig Holstein, Kiel 24105, Germany.

51. Department of Neurology, NYU School of Medicine, New York City, NY 10003, USA.

52. Department of Neurology, Charité Universitaetsmedizin Berlin, Campus Virchow-Clinic, Berlin 13353, Germany.

53. Institute of Neurology and Neurosurgery at St. Barnabas, Livingston, NJ 07039, USA.

54. Research Unit of Molecular Epidemiology, Helmholtz Zentrum München - German Research Center for Environmental Health, Neuherberg D-85764, Germany.

55. Institute of Epidemiology, Helmholtz Zentrum München - German Research Center for Environmental Health, Neuherberg D-85764, Germany.

56. Comprehensive Epilepsy Center, Division of Neurology, Cincinnati Children's Hospital Medical Center, Cincinnati, OH 45229, USA.

57. Department of Neurology, University of Michigan, Ann Arbor, MI 48109, USA.

58. Center for Human Genome Variation, Duke University School of Medicine, Durham, NC 27710, USA.

59. Institute for Genomic Medicine, Columbia University Medical Center, New York, NY 10032, USA.

60. Division of Human Genetics, Department of Pediatrics, The Perelman School of Medicine, University of Pennsylvania, Philadelphia, PA 19104, USA.

61. Life and Brain Center, University of Bonn Medical Center, Bonn 53127, Germany.

62. Montefiore Medical Center, Bronx, NY 10467, USA.

63. Department of Neuropediatrics, University Medical Center Schleswig-Holstein (UKSH), Kiel 24105, Germany.

64. Danish Epilepsy Centre, Dianalund 4293, Denmark.

65. Institute of Regional Health Services Research, University of Southern Denmark, Odense 5000, Denmark.

66. deCODE genetics, Reykjavik IS-101, Iceland.

67. Department of Psychiatry and Applied Psychology, Institute of Mental Health University of Nottingham, Nottingham NG7 2TU, UK.

68. Faculty of Medicine, Imperial College London, London SW7 2AZ, UK.

69. Kuopio Epilepsy Center, Neurocenter, Kuopio University Hospital, Kuopio 70029, Finland.

70. Institute of Clinical Medicine, University of Eastern Finland, Kuopio 70029, Finland.

71. Department of Neurology, University of California, San Francisco, CA 94143, USA.

72. University of Alabama Birmingham, Department of Neurology, Birmingham, AL 35233, USA.

73. Luxembourg Centre for Systems Biomedicine, University of Luxembourg, Esch-sur-Alzette L-4362, Luxembourg.

74. Department of Neurology, Medical University of Vienna, Vienna 1090, Austria.

75. Department of Neurology, Zucker-Hofstra Northwell School of Medicine, NY 10075, USA.

76. Department of Medicine, University of Melbourne, Royal Melbourne Hospital, Parkville 3050, Australia.

77. Department of Neuroscience, Central Clinical School, Monash University, Melbourne 3004, Australia.

78. Stanley Center for Psychiatric Research, Broad Institute of Harvard and M.I.T., Cambridge, MA 02142, USA.

79. Department of Paediatrics and Adolescent Medicine, The University of Hong Kong, Hong Kong.

80. Folkhälsan Research Center and Medical Faculty, University of Helsinki, Helsinki 00290, Finland.

81. Genomic Medicine Institute, Lerner Research Institute, Cleveland Clinic, Cleveland, OH 44195, USA.

82. Institut für Epidemiologie Christian-Albrechts-Universität zu Kiel, Kiel 24105, Germany.

83. Department of Pediatrics and Neurology, Ohio State University and Nationwide Children's Hospital, Columbus, OH 43205, USA.

84. Department of Medical Genetics, School of Medical Sciences, University of Campinas (UNICAMP), Campinas, SP 13083-887, Brazil.

85. Brazilian Institute of Neuroscience and Neurotechnology (BRAINN), Campinas, SP 13083-970, Brazil.

86. Istituti Clinici Scientifici Maugeri, Pavia 27100, Italy.

87. Epilepsy Center Kleinwachau, Radeberg 01454, Germany.

88. Division of Intramural Population Health Research, Eunice Kennedy Shriver National Institute of Child Health and Human Development, National Institutes of Health, Bethesda, MD 20892, USA*.*

89. Wilhelm Johannsen Centre for Functional Genome Research, Copenhagen DK-2200, Denmark.

90. School of Medicine, Trinity College Dublin, Dublin 2, Ireland.

91. Department of Neurology, Austin Health, Heidelberg 3084, Australia.

92. United Christian Hospital, Hong Kong.

93. Institute of Human Genetics, University of Bonn Medical Center, Bonn 53127, Germany.

94. Cologne Center for Genomics, University of Cologne, Cologne 50931, Germany.

95. Institute for Molecular Medicine Finland (FIMM), University of Helsinki, Helsinki 0014, Finland*.*

96. The Broad Institute of M.I.T. and Harvard, Cambridge, MA 02142, USA.

97. AstraZeneca Centre for Genomics Research, Precision Medicine and Genomics, IMED Biotech Unit, AstraZeneca, Cambridge CB2 0AA, UK.

98. Department of Neurology, Boston Children's Hospital, Harvard Medical School, Boston, MA 02115, USA.

99. Department of Neurology, Neuroscience Institute, University of Cincinnati Medical Center, Cincinnati, OH 45220, USA.

100. Department of Neurology, Duke University School of Medicine, Durham, NC 27710, USA.

101. Epilepsy-Center Hessen, Department of Neurology, University Medical Center Giessen and Marburg, Marburg, Germany and Philipps-University Marburg, Marburg 35043, Germany.

102. Epilepsy Center Frankfurt Rhine-Main, Center of Neurology and Neurosurgery, Goethe University Frankfurt, Frankfurt 60528, Germany.

103. Chalfont Centre for Epilepsy, Chalfont-St-Peter, Buckinghamshire SL9 0RJ, UK.

104. Department of Endocrinology, Hospital of The University of Pennsylvania, Philadelphia, PA 19104, USA.

105. Departments of Neurology, Beth Israel Deaconess Medical Center, Massachusetts General Hospital, and Harvard Medical School, Boston, MA 02215, USA.

106. Department of Neurology, Inselspital, Bern University Hospital, University of Bern, Bern 3010, Switzerland.

107. Department of Neurology, Royal Children's Hospital, Parkville 3052, Australia.

108. Department of Neurosciences, University of California, San Diego, La Jolla, CA 92037, USA.

109. The Royal College of Surgeons in Ireland, Dublin D02 YN77, Ireland. *.*

110. Rush University Medical Center, Chicago, IL 60612, USA.

111. Department of Neurology, Alan Richens Epilepsy Unit, University Hospital of Wales, Cardiff CF14 4XW, UK.

112. Aarhus Institute of Advanced Studies (AIAS), Aarhus University, 8000 Aarhus, Denmark.

113. Department of Neurology and Comprehensive Epilepsy Center, Thomas Jefferson University, Philadelphia, PA 19107, USA.

114. Institute of Genetic Epidemiology, Helmholtz Zentrum München - German Research Center for Environmental Health, Neuherberg D-85764, Germany.

115. Chair of Genetic Epidemiology, IBE, Faculty of Medicine, LMU Munich 80539, Germany.

116. Pediatric Neurology and Muscular Diseases Unit, Department of Neurosciences, Rehabilitation, Ophthalmology, Genetics, Maternal and Child Health, G. Gaslini Institute, University of Genoa, Genova 16148, Italy.

117. CWZ Hospital, 6532 SZ Nijmegen, The Netherlands.

118. Department of Neurology, Washington University School of Medicine, St. Louis, MO 63110, USA*.*

119. Institute for Applied Health Research, University of Birmingham, Birmingham B15 2TT, UK.

120. C. Mondino National Neurological Institute, Pavia 27100, Italy.

121. Departments of Neurology and Pediatrics, The Johns Hopkins University School of Medicine, Baltimore, MD 21287, USA.

122. Department of Neurology, Admiraal De Ruyter Hospital, Goes 4462, The Netherlands.

123. Division of Medical Genetics, Department of Pediatrics, Duke University Medical Center, Durham, NC 27710, USA.

124. Department of Computer Science, New Jersey Institute of Technology, NJ 07102, USA.

125. Department of Pediatric Neurology and Developmental Medicine, University Children's Hospital, Tübingen 72076, Germany.

126. Department of Neurology, Institute of Clinical Medicine, Faculty of Medicine, Vilnius University, Vilnius, Lithuania.

**1. Epi25 Consortium**

**Epi25 sequencing, analysis, and project management at the Broad Institute**

Yen-Chen Anne Feng^1-4^, Daniel P. Howrigan^1,3,4^, Liam E. Abbott^1,3,4^, Katherine Tashman^1,3,4^, Felecia Cerrato^3^, Dennis Lal^4,5^, Costin Leu^4,5^, Claire Churchhouse^1,3,4^, Namrata Gupta^3^, Stacey B. Gabriel^146^, Mark J. Daly^1,3,4^, Eric S. Lander^146,147,148^, Benjamin M. Neale^1,3,4^

**Epi25 executive committee**

Samuel F. Berkovic ^9^, Holger Lerche^8^, David B. Goldstein ^6^, Daniel H. Lowenstein^7^

**Epi25 strategy, phenotyping, analysis, informatics, and project management committees**

Samuel F. Berkovic^9^, Holger Lerche^8^, David B. Goldstein^6^, Daniel H. Lowenstein^7^, Gianpiero L. Cavalleri^67,70^, Patrick Cossette^106^, Chris Cotsapas^111^, Peter De Jonghe^12-14^, Tracy Dixon-Salazar^112^, Renzo Guerrini^82^, Hakon Hakonarson^101^, Erin L. Heinzen^6^, Ingo Helbig^28,29,102^, Patrick Kwan^10,11^, Anthony G. Marson^58^, Slavé Petrovski^11,116^, Sitharthan Kamalakaran^6^, Sanjay M. Sisodiya^57^, Randy Stewart^113^, Sarah Weckhuysen^12-14^, Chantal Depondt^15^, Dennis J. Dlugos^101^, Ingrid E. Scheffer^9^, Pasquale Striano^74^, Catharine Freyer^7^, Roland Krause^114^, Patrick May^114^, Kevin McKenna^7^, Brigid M. Regan^9^, Susannah T. Bellows^9^, Costin Leu^4,5,57^

**Authors from individual Epi25 cohorts:**

**Australia: Melbourne (AUSAUS)**

Samuel F. Berkovic^9^, Ingrid E. Scheffer^9^, Brigid M. Regan^9^, Caitlin A. Bennett^9^, Susannah T. Bellows^9^, Esther M.C. Johns^9^, Alexandra Macdonald^9^, Hannah Shilling^9^, Rosemary Burgess^9^,

Dorien Weckhuysen^9^, Melanie Bahlo^119,120^

**Australia: Royal Melbourne (AUSRMB)**

Terence J. O'Brien^10,11^, Patrick Kwan^10,11^, Slavé Petrovski^11,116^, Marian Todaro^10,11^

**Belgium: Antwerp (BELATW)**

Sarah Weckhuysen^12-14^, Hannah Stamberger^12-14^, Peter De Jonghe^12-14^

**Belgium: Brussels (BELULB)**

Chantal Depondt^15^

**Canada: Andrade (CANUTN)**

Danielle M. Andrade^16,17^, Tara R. Sadoway^17^, Kelly Mo^17^

**Switzerland: Bern (CHEUBB)**

Heinz Krestel^18^, Sabina Gallati^19^

**Cyprus (CYPCYP)**

Savvas S. Papacostas^20^, Ioanna Kousiappa^20^, George A. Tanteles^21^

**Czech Republic: Prague (CZEMTH)**

Katalin Štěrbová^22^, Markéta Vlčková^23^, Lucie Sedláčková^22^, Petra Laššuthová^22^

**Germany: Frankfurt/Marburg (DEUPUM)**

Karl Martin Klein^24,25^, Felix Rosenow^24,25^, Philipp S. Reif^24,25^, Susanne Knake^25^

**Germany: Bonn (DEUUKB)**

Wolfram S. Kunz^26,27^, Gábor Zsurka^26,27^, Christian E. Elger^27^, Jürgen Bauer^27^, Michael Rademacher^27^

**Germany: Kiel (DEUUKL)**

Ingo Helbig^28,29,102^, Karl Martin Klein^24,25^, Manuela Pendziwiat^29^, Hiltrud Muhle^29^, Annika Rademacher^29^, Andreas van Baalen^29^, Sarah von Spiczak^29^, Ulrich Stephani^29^, Zaid Afawi^30^, Amos D. Korczyn^31^, Moien Kanaan^32^, Christina Canavati^32^, Gerhard Kurlemann^33^, Karen Müller-Schlüter^34^, Gerhard Kluger^35,36^, Martin Häusler^37^, Ilan Blatt^31,115^

**Germany: Leipzig (DEUULG)**

Johannes R. Lemke^38^, Ilona Krey^38^

**Germany: Tuebingen (DEUUTB)**

Holger Lerche^8^, Yvonne G. Weber^8,151^, Stefan Wolking^8^, Felicitas Becker^8,39^, Christian Hengsbach^8^, Sarah Rau^8^, Ana F. Maisch^8^, Bernhard J. Steinhoff^40^, Andreas Schulze-Bonhage^41^, Susanne Schubert-Bast^42^, Herbert Schreiber^43^, Ingo Borggräfe^44^, Christoph J. Schankin^45^, Thomas Mayer^46^, Rudolf Korinthenberg^47^, Knut Brockmann^48^, Gerhard Kurlemann^33^, Dieter Dennig^49^, Rene Madeleyn^50^

**Finland: Kuopio (FINKPH)**

Reetta Kälviäinen^51^, Pia Auvinen^51^, Anni Saarela^51^

**Finland: Helsinki (FINUVH)**

Tarja Linnankivi^52^, Anna-Elina Lehesjoki^53^

**Wales: Swansea (GBRSWU)**

Mark I. Rees^54,55^, Seo-Kyung Chung^54,55^, William O. Pickrell^54^, Robert Powell^54,56^

**UK: UCL (GBRUCL)**

Sanjay M. Sisodiya^57^, Natascha Schneider^57^, Simona Balestrini^57^, Sara Zagaglia^57^, Vera Braatz^57^

**UK: Imperial/Liverpool (GBRUNL)**

Anthony G. Marson^58^, Michael R. Johnson^59^, Pauls Auce^60^, Graeme J. Sills^61^

**Hong Kong (HKGHKK)**

Patrick Kwan^10,11,62^, Larry W. Baum^117,118,63^, Pak C. Sham^117,118,63^, Stacey S. Cherny^64^, Colin H.T. Lui^65^

**Croatia (HRVUZG)**

Nina Barišić^66^

**Ireland: Dublin (IRLRCI)**

Gianpiero L. Cavalleri^67,70^, Norman Delanty^67,70^, Colin P. Doherty^68,70^, Arif Shukralla^69^, Mark McCormack^67^, Hany El-Naggar^69,70^

**Italy: Milan (ITAICB)**

Laura Canafoglia^71^, Silvana Franceschetti^71^, Barbara Castellotti^72^, Tiziana Granata^73^

**Italy: Genova	(ITAIGI)**

Pasquale Striano^74^, Federico Zara^75^, Michele Iacomino^75^, Francesca Madia^75^, Maria Stella Vari^74^, Maria Margherita Mancardi^75^, Vincenzo Salpietro^74^

**Italy: Bologna (ITAUBG)**

Francesca Bisulli^76,77^, Paolo Tinuper^76,77^, Laura Licchetta^76,77^, Tommaso Pippucci^78^, Carlotta Stipa^79^, Raffaella Minardi^76^

**Italy: Catanzaro (ITAUMC)**

Antonio Gambardella^80^, Angelo Labate^80^, Grazia Annesi^81^, Lorella Manna^81^, Monica Gagliardi^81^

**Italy: Florence (ITAUMR)**

Renzo Guerrini^82^, Elena Parrini^82^, Davide Mei^82^, Annalisa Vetro^82^, Claudia Bianchini^82^, Martino Montomoli^82^, Viola Doccini^82^, Carla Marini^82^

**Japan: RIKEN Institute (JPNRKI)**

Toshimitsu Suzuki^83^, Yushi Inoue^84^, Kazuhiro Yamakawa^83^

**Lithuania (LTUUHK)**

Birute Tumiene^85,86^

**New Zealand: Otago (NZLUTO)**

Lynette G. Sadleir^87^, Chontelle King^87^, Emily Mountier^87^

**Turkey: Bogazici (TURBZU)**

S. Hande Caglayan^88^, Mutluay Arslan^89^, Zuhal Yapıcı^90^, Uluc Yis^91^, Pınar Topaloglu^90^, Bulent Kara^92^, Dilsad Turkdogan^93^, Aslı Gundogdu-Eken^88^

**Turkey: Istanbul (TURIBU)**

Nerses Bebek^94,95^, Sibel Uğur-İşeri^95^, Betül Baykan^94^, Barış Salman^95^, Garen Haryanyan^94^, Emrah Yücesan^149^, Yeşim Kesim^94^, Çiğdem Özkara^96^

**USA: BCH (USABCH)**

Annapurna Poduri^97,98^

**USA: Philadelphia/CHOP (USACHP) and Philadelphia/Rowan (USACRW)**

Russell J. Buono^99,100,101^, Thomas N. Ferraro^99,102^, Michael R. Sperling^100^, Dennis J. Dlugos^101,102^, Warren Lo^103^, Michael Privitera^104^, Jacqueline A. French^105^, Patrick Cossette^106^, Steven Schachter^107^, Hakon Hakonarson^101^

**USA: EPGP (USAEGP)**

Daniel H. Lowenstein^7^, Ruben I. Kuzniecky^108^, Dennis J. Dlugos^101,102^, Orrin Devinsky^105^

**USA: NYU HEP (USAHEP)**

Daniel H. Lowenstein^7^, Ruben I. Kuzniecky^108^, Jacqueline A. French^105^, Manu Hegde^7^

**USA: Penn/CHOP (USAUPN)**

Ingo Helbig^28,102^, Pouya Khankhanian^109,110^, Katherine L. Helbig^28^, Colin A. Ellis^110^

Affiliations:

1 Analytic and Translational Genetics Unit, Department of Medicine, Massachusetts General Hospital and Harvard Medical School, Boston, MA 02114, USA
2 Psychiatric & Neurodevelopmental Genetics Unit, Department of Psychiatry, Massachusetts General Hospital and Harvard Medical School, Boston, MA 02114, USA
3 Program in Medical and Population Genetics, Broad Institute of Harvard and MIT, 7 Cambridge Center, Cambridge, MA 02142, USA
4 Stanley Center for Psychiatric Research, Broad Institute of Harvard and MIT, Cambridge, MA 02142, USA
5 Genomic Medicine Institute, Cleveland Clinic, Cleveland, OH 44195, USA
6 Institute for Genomic Medicine, Columbia University, New York, NY 10032, USA
7 Department of Neurology, University of California, San Francisco, CA 94110, USA
8 Department of Neurology and Epileptology, Hertie Institute for Clinical Brain Research, University of Tübingen, 72076 Tübingen, Germany
9 Epilepsy Research Centre, Department of Medicine, University of Melbourne, Victoria, Australia
10 Department of Neuroscience, Central Clinical School, Monash University, Alfred Hospital, Melbourne, Australia
11 Departments of Medicine and Neurology, University of Melbourne, Royal Melbourne Hospital, Parkville, Australia
12 Neurogenetics Group, Center for Molecular Neurology, VIB, Antwerp, Belgium
13 Laboratory of Neurogenetics, Institute Born-Bunge, University of Antwerp, Belgium
14 Division of Neurology, Antwerp University Hospital, Antwerp, Belgium
15 Department of Neurology, Université Libre de Bruxelles, Brussels, Belgium
16 Department of Neurology, Toronto Western Hospital, Toronto, ON M5T 2S8, Canada
17 University Health Network, University of Toronto, Toronto, ON, Canada
18 Departments of Neurology and BioMedical Research, Bern University Hospital and University of Bern, Bern, Switzerland
19 Institute of Human Genetics, Bern University Hospital, Bern, Switzerland
20 Neurology Clinic B, The Cyprus Institute of Neurology and Genetics, 2370 Nicosia, Cyprus
21 Department of Clinical Genetics, The Cyprus Institute of Neurology and Genetics, 2370 Nicosia, Cyprus
22 Department of Paediatric Neurology, 2nd Faculty of Medicine, Charles University and Motol Hospital, Prague, Czech Republic
23 Department of Biology and Medical Genetics, 2nd Faculty of Medicine, Charles University and Motol Hospital, Prague, Czech Republic
24 Epilepsy Center Frankfurt Rhine-Main, Center of Neurology and Neurosurgery, Goethe University Frankfurt, Frankfurt, Germany
25 Epilepsy Center Hessen-Marburg, Department of Neurology, Philipps University Marburg, Marburg, Germany
26 Institute of Experimental Epileptology and Cognition Research, University Bonn, 53127

Bonn, Germany

27 Department of Epileptology, University Bonn, 53127 Bonn, Germany
28 Division of Neurology, Children's Hospital of Philadelphia, Philadelphia, PA 19104, USA
29 Department of Neuropediatrics, Christian-Albrechts-University of Kiel, 24105 Kiel, Germany
30 Sackler School of Medicine, Tel-Aviv University, Ramat Aviv, Israel
31 Tel-Aviv University Sackler Faculty of Medicine, Ramat Aviv 69978, Israel
32 Hereditary Research Lab, Bethlehem University, Bethlehem, Palestine
33 Department of Neuropediatrics, Westfälische Wilhelms-University, Münster, Germany
34 Epilepsy Center for Children, University Hospital Neuruppin, Brandenburg Medical School, Neuruppin, Germany
35 Neuropediatric Clinic and Clinic for Neurorehabilitation, Epilepsy Center for Children and Adolescents, Vogtareuth, Germany
36 Research Institute Rehabilitation / Transition / Palliation, PMU Salzburg, Austria
37 Division of Neuropediatrics and Social Pediatrics, Department of Pediatrics, University Hospital, RWTH Aachen, Aachen, Germany
38 Institute of Human Genetics, Leipzig, Germany
39 RKU-University Neurology Clinic of Ulm, Ulm, Germany
40 Kork Epilepsy Center, Kehl-Kork, Germany
41 Epilepsy Center, University of Freiburg, Freiburg im Breisgau, Germany
42 Section Neuropediatrics and Inborn Errors of Metabolism, University Children's Hospital, Heidelberg, Germany
43 Neurological Practice Center & Neuropoint Patient Academy, Ulm, Germany.
44 Department of Pediatric Neurology and Developmental Medicine, LMU Munich, Munich, Germany
45 Department of Neurology, University of Munich Hospital-Großhadern, Munich, Germany
46 Saxonian Epilepsy Center Radeberg, Radeberg, Germany
47 Division of Neuropediatrics and Muscular Disorders, University Hospital Freiburg, Freiburg, Germany
48 University Children's Hospital, Göttingen, Germany
49 Private Neurological Practice, Stuttgart, Germany
50 Department of Pediatrics, Filderklinik, Filderstadt, Germany
51 Neurocenter, Kuopio University Hospital, Kuopio Finland and Institute of Clinical Medicine, University of Eastern Finland, Finland
52 Child Neurology, University of Helsinki and Helsinki University Hospital, Helsinki, Finland
53 Medicum, University of Helsinki, Helsinki, Finland and Folkhälsan Research Center, Helsinki, Finland
54 Neurology Research Group, Swansea University Medical School, Swansea University SA2 8PP, UK
55 Faculty of Medicine and Health, University of Sydney, Sydney, Australia
56 Department of Neurology, Morriston Hospital, Abertawe BroMorgannwg HealthBoard, Swansea, UK
57 Department of Clinical and Experimental Epilepsy, UCL Queen Square Institute of Neurology, London, UK and Chalfont Centre for Epilepsy, Chalfont St Peter, UK
58 Department of Molecular and Clinical Pharmacology, University of Liverpool, Liverpool, UK
59 Division of Brain Sciences, Imperial College London, London, UK
60 Department of Neurology, Walton Centre NHS Foundation Trust, Liverpool, UK
61 School of Life Sciences, University of Glasgow, Glasgow, UK
62 Department of Medicine and Therapeutics, Chinese University of Hong Kong, Hong Kong, China
63 Department of Psychiatry, University of Hong Kong, Hong Kong, China
64 Department of Epidemiology and Preventive Medicine and Department of Anatomy and Anthropology, Sackler Faculty of Medicine, Tel Aviv University, Israel
65 Department of Medicine, Tseung Kwan O Hospital, Hong Kong, China
66 Department of Pediatric University Hospital centre Zagreb, Croatia
67 The Department of Molecular and Cellular Therapeutics, The Royal College of Surgeons in Ireland, Dublin, Ireland
68 Neurology Department, St. James Hospital, Dublin, Ireland
69 The Department of Neurology, Beaumont Hospital, Dublin, Ireland
70 The FutureNeuro Research Centre, Ireland
71 Neurophysiopathology, Fondazione IRCCS Istituto Neurologico Carlo Besta, Milan, Italy
72 Unit of Genetics of Neurodegenerative and Metabolic Diseases, Fondazione IRCCS Istituto Neurologico Carlo Besta, Milan, Italy
73 Department of Pediatric Neuroscience, Fondazione IRCCS Istituto Neurologico Carlo Besta, Milan, Italy
74 Pediatric Neurology and Muscular Diseases Unit, Department of Neurosciences, Rehabilitation, Ophthalmology, Genetics, Maternal and Child Health, University of Genoa, "G. Gaslini" Institute, Genova, Italy
75 Laboratory of Neurogenetics, "G. Gaslini" Institute, Genova, Italy
76 IRCCS, Institute of Neurological Sciences of Bologna, Bologna, Italy
77 Department of Biomedical and Neuromotor Sciences, University of Bologna, Bologna, Italy
78 Medical Genetics Unit, Polyclinic Sant'Orsola-Malpighi University Hospital, Bologna, Italy
79 Department of Biomedical and Neuromotor Sciences, University of Bologna, Bologna, Italy
80 Institute of Neurology, Department of Medical and Surgical Sciences, University “Magna Graecia”, Catanzaro, Italy
81 Institute of Molecular Bioimaging and Physiology, CNR, Section of Germaneto, Catanzaro, Italy
82 Pediatric Neurology, Neurogenetics and Neurobiology Unit and Laboratories, Children's Hospital A. Meyer, University of Florence, Italy
83 Laboratory for Neurogenetics, RIKEN Center for Brain Science, Saitama, Japan
84 National Epilepsy Center, Shizuoka Institute of Epilepsy and Neurological Disorder, Shizuoka, Japan
85 Institute of Biomedical Sciences, Faculty of Medicine, Vilnius University, Vilnius, Lithuania
86 Centre for Medical Genetics, Vilnius University Hospital Santaros Klinikos, Vilnius, Lithuania
87 Department of Paediatrics and Child Health, University of Otago, Wellington
88 Department of Molecular Biology and Genetics, Bogaziçi University, Istanbul, Turkey
89 Department of Child Neurology, Gulhane Education and Research Hospital, Health Sciences University, Ankara, Turkey
90 Department of Child Neurology, Istanbul Faculty of Medicine, Istanbul University, Istanbul, Turkey
91 Department of Child Neurology, Medical School, Dokuz Eylul University, Izmir, Turkey
92 Department of Child Neurology, Medical School, Kocaeli University, Kocaeli, Turkey
93 Department of Child Neurology, Medical School, Marmara University, Istanbul, Turkey
94 Department of Neurology, Istanbul Faculty of Medicine, Istanbul University, Istanbul, Turkey
95 Department of Genetics, Aziz Sancar Institute of Experimental Medicine, Istanbul University, Istanbul, Turkey
96 Department of Neurology, Faculty of Medicine, Cerrahpaşa University Istanbul, Istanbul, Turkey
97 Epilepsy Genetics Program, Department of Neurology, Boston Children's Hospital, Boston, MA 02115, USA
98 Department of Neurology, Harvard Medical School, Boston, MA 02115, USA
99 Cooper Medical School of Rowan University, Camden, NJ 08103, USA
100 Thomas Jefferson University, Philadelphia, PA 19107, USA
101 The Children's Hospital of Philadelphia, Philadelphia, PA 19104, USA
102 Perelman School of Medicine, University of Pennsylvania, PA 19104, USA
103 Nationwide Children's Hospital, Columbus, OH 43205, USA
104 University of Cincinnati, Cincinnati, OH 45220, USA
105 Department of Neurology, New York University/Langone Health, New York, NY 10016, USA
106 University of Montreal, Montreal, QC H3T 1J4, Canada
107 Beth Israel Deaconess/Harvard, Boston, MA 02115, USA
108 Department of Neurology, Hofstra-Northwell Medical School, New York, NY 11549, USA
109 Center for Neuro-engineering and Therapeutics, University of Pennsylvania, Philadelphia, PA 19104

, USA

110 Department of Neurology, Hospital of University of Pennsylvania, Philadelphia, PA 19104, USA
111 School of Medicine, Yale University, New Haven, CT 06510, USA
112 LGS Foundation, NY 11716, USA
113 National Institute of Neurological Disorders and Stroke, MD 20852, USA
114 Luxembourg Centre for Systems Biomedicine, University Luxembourg, Esch-sur-Alzette, Luxembourg
115 Department of Neurology, Sheba Medical Center, Ramat Gan, Israel
116 Centre for Genomics Research, Precision Medicine and Genomics, IMED Biotech Unit, AstraZeneca, Cambridge, UK
117 The State Key Laboratory of Brain and Cognitive Sciences, University of Hong Kong, Hong Kong, China
118 Centre for Genomic Sciences, University of Hong Kong, Hong Kong, China
119 Population Health and Immunity Division, the Walter and Eliza Hall Institute of Medical Research, Parkville 3052, VIC, Australia
120 Department of Medical Biology, The University of Melbourne, Melbourne 3010, VIC, Australia
